# A high-quality genome assembly of the ghost moth *Druceiella hillmani* provides new evidence of genome size augmentation in Hepialidae

**DOI:** 10.1101/2023.12.05.570119

**Authors:** Yi-Ming Weng, Isabel Lopez-Cacacho, Bert Foquet, Jose I. Martinez, David Plotkin, Andrei Sourakov, Akito Y. Kawahara

**Author notes:** Corresponding author: Akito Y. Kawahara.

## Abstract

Ghost moths are an unusual family of primitive moths (Lepidoptera: Hepialidae) known for their large body size and crepuscular adult activity. These moths represent an ancient lineage, frequently have soil dwelling larvae, and are adapted to high elevations, deserts, and other extreme environments. Despite being rather speciose with more than 700 species, there is a dearth of genomic resources for the family. Here, we present the first high quality, publicly available hepialid genome, generated from an Andean species of ghost moth, *Druceiella hillmani*. Our genome assembly has a length of 2,586 Mbp with contig N50 of 28.1 Mb and N50 of 29, and BUSCO completeness of 97.1%, making it one of the largest genomes in the order Lepidoptera. Our assembly is a vital resource for future research on ghost moth genomics.

## Background & Summary

Lepidoptera (butterflies and moths) are one of the most widely studied arthropod orders, with over 160,000 described species. Most of this work focuses on Lepidoptera in the megadiverse clade Ditrysia, with the non-ditrysian moths (representing < 2% of all Lepidoptera species) receiving much less attention^1^. One of these non-ditrysian lineages is Hepialidae, an enigmatic family commonly known as ghost moths or swift moths. Hepialidae comprise roughly 700 species and are found on all continents except Antarctica, with most species adapted to living at high elevations and other extreme environments. Hepialid adults have specialized genitalic structures, and during reproduction they exhibit an unusual method of external sperm transfer^2,3^. Most ghost moth caterpillars are subterranean, feeding on roots or mycelia, and creating underground silk tunnels when moving to new host plants and host fungi^4^. Hepialidae are thought to have the largest genomes of any Lepidoptera, but the causes and implications of this genome expansion have not been extensively investigated due to an absence of high-quality hepialid genomic data^5,6^.

Here, we sequenced and assembled the genome of the ghost moth *Druceiella hillmani*, a neotropical species that lives at high elevations in Ecuadorian regions of the Andes^7^. The genome size of *D. hillmani* turned out to be significantly larger than that of most other lepidopteran families, almost doubling the genome assembly size of *Parnassius apollo*, which was considered the largest assembled genome at the time^8^. Our genome assembly supports the previous hypotheses about the large genomes in this family. Our study provides the first high-quality, publicly available genome assembly for the Hepialidae family, and will serve as a foundation for future research on the evolution of the ghost moths’ fascinating reproductive and ecological adaptations.

## Methods

### Sampling and Sequencing

We collected an adult female ghost moth, *Druceiella hillmani* (Hepialidae), from WildSumaco Biological Station, Napo Province, Ecuador (0°40’57”S 77°35’35”W, 1400 m) on June 4, 2023, with permit number MAATE-ARSFC-2023-3144, and deposited it in the collection of the McGuire Center for Lepidoptera and Biodiversity at the Florida Museum of Natural History (FLMNH MGCL Wing Voucher Accession Number: LEP-89533) (**Figure 1**). Genomic DNA was extracted from whole body tissue (except for the abdomen) using a Qiagen Genomic-tip DNA extraction kit (Qiagen, LLC). Genomic libraries were constructed using the SMRTbell Express Template Prep Kit 2.0 (PacBio, Menlo Park, CA, USA). PacBio HiFi whole-genome sequencing in circular consensus sequencing (ccs) mode was performed using a PacBio Sequel II instrument at the DNA Sequencing Center at Brigham Young University (Provo, Utah, USA).

**Figure 1.**
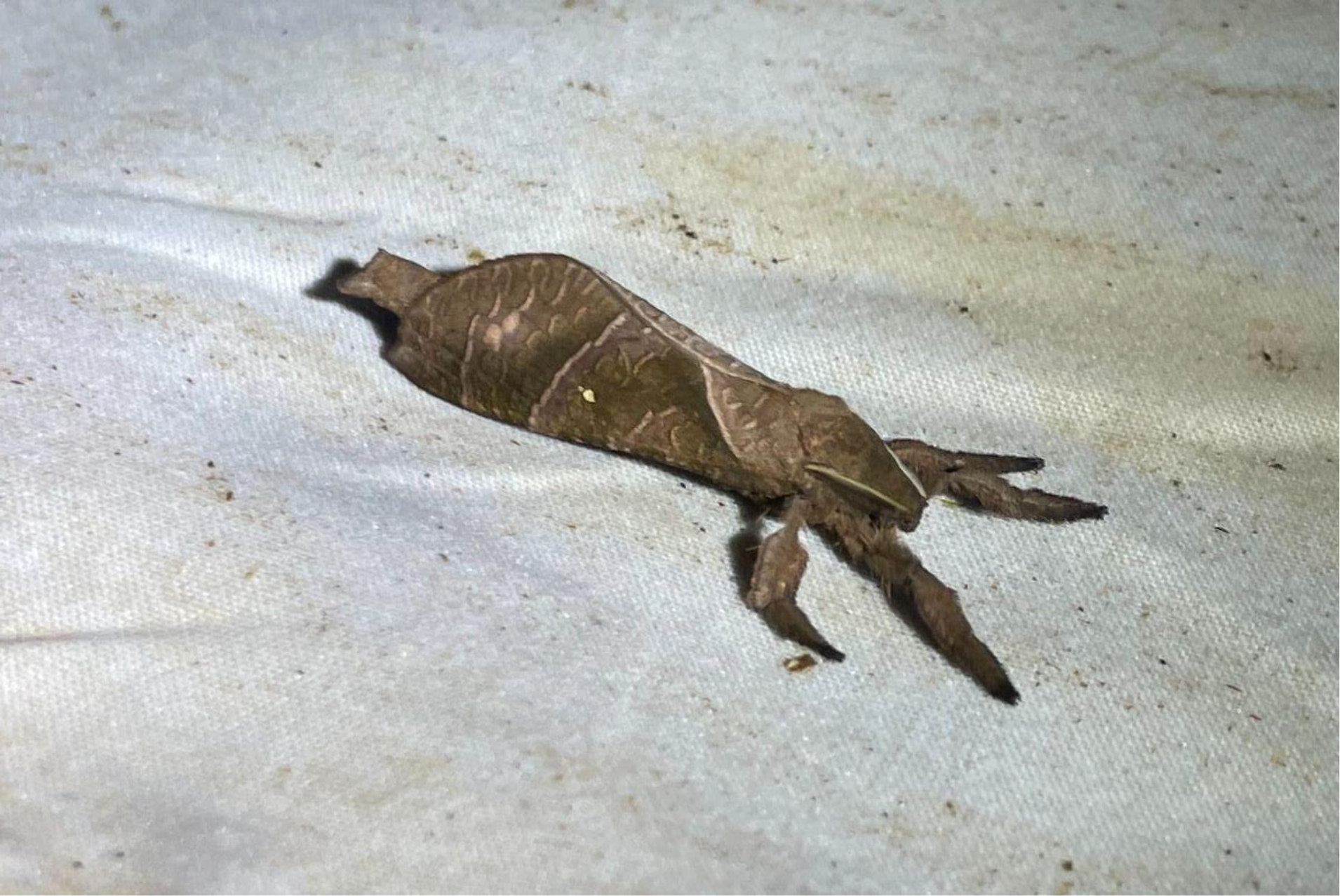
Female specimen of *Druceiella hillmani* (Hepialidae) that was sampled to construct the genome in the present study.

### Genome assembly and quality assessment

We assembled the genome using our previously published genome-assembly pipeline^9^. Briefly, raw HiFi ccs reads were fed to the Hifiasm assembler and the assembled output was purged for duplicated haplotigs using purge_haplotigs v1.1.2^10,11^. In this pipeline, minimap2 was used to map the ccs reads to the raw assembly and the read depth at low and high peaks as well as the lowest midpoint were assigned to 5, 100, and 17, respectively, according to the mapped coverage histogram^12^ (**Supplementary Figure S1**). Non-target DNA detection was performed using blobtools^13^, where the top hit from the megablast result with an e-value cutoff of 1E-25 was used to assign the taxonomic ID to the contigs. The contig removal was based on the deviations of GC content, mean sequence coverage, and the assigned taxon (**Supplementary Figure S2**). The assembly completeness was accessed using QUAST v5.0.2 and BUSCO v5.3.0, based on the endopterygota_odb10 database^14,15^. Finally we assigned the assembled contigs to putative chromosomes by linking the BUSCO single-copy genes to the reference genome of one other moth species, *Tinea pellionella* (accession: GCA_948150575) from an ancient derived lineage, based on the fact that most Lepidoptera species have holocentric chromosomes with high chromosome-level gene collinearity^16^. Specifically, we downloaded the high-quality chromosome-level assembly of *T. pellionella* from the Darwin Tree of Life genome database (DToL, https://portal.darwintreeoflife.org/) and performed BUSCO analysis on this genome with the same settings as described above. The chromosomal coordinates for each single copy ortholog found in both species were extracted from the *T. pellionella* assembly, and these coordinates were used to link each contig of the *D. hillmani* genome to the chromosome with which it shares the highest number of BUSCO orthologs (**Supplementary Data1**).

### Genome annotation

Feature and functional annotation was performed on the final assembly, after duplicated haplotig purging and non-target DNA removal steps. We first used RepeatModeler2 to build a *de novo* repeat library which was then used together with the repeat database of Arthropoda from Repbase to soft-mask the genome using RepeatMasker^17–19^. Since Helitrons (a superfamily of transposons) have been found to be particularly more prevalent in Lepidoptera, and have been shown to influence the composition of the host genome^20,21^, we scanned for Helitron-related sequences using HelitronScanner with default settings^22^. In addition, we searched for non-coding RNAs (ncRNA) using Infernal where the RNA database from Rfam was used as reference to further explore the genome constitution of this genome assembly^23,24^. The soft-masked genome from RepeatMasker was subsequently used to predict coding genes using fully automatic BRAKER3 gene annotation pipeline. The arthropod protein sequences from OrthoDB v11 were used to generate hints in the ProtHint pipeline for training GeneMark-EP+ and Augustus gene prediction^25–33^. The gene model was summarized using gFACs v1.1.2 and the gene model completeness was assessed using BUSCO v5.3.0 based on the endopterygota_odb10 database^14,34^. The Augustus gene model predicted from the BRAKER3 pipeline was then used to perform the functional annotation, using eggNOG-mapper v2.1.6 with default settings and DIAMOND v2.0.9 to blast sequences against the RefSeq non-redundant (nr) protein database with an e-value cutoff of 1E-10^30,35,36^. From the blast results, the best hit with the lowest e-value was selected and the corresponding species and its order were extracted. We mapped the order of the genes to the contigs for a visual examination of potential non-target DNA from parasitoids and gut contents.

### Genome size estimation

The genome size of *D. hillmani* was estimated using two different approaches. First, we used KMC to count the k-mers with size of 21bps^37^. The k-mer count distribution was then fed to GenomeScope2 to estimate the genome size, sequence coverage, and heterozygosity with default settings^38^. Second, we used backmap (ModEst) to map the raw ccs reads to the final assembly to again estimate the sequence coverage and genome size^12,39–43^. Both estimates were compared with the final assembly size.

To compare genome sizes across Lepidoptera species, we retrieved 2,079 assemblies from 961 lepidopteran species from the Arthropoda Assembly Assessment Catalogue (A^3^Cat, https://a3cat.unil.ch, accessed date: November 21^th^ 2023). Assemblies were selected based on their BUSCO scores: only assemblies with a complete score > 80% and duplication rate < 5% were included for the analysis. These filters ensured the completeness and cleanness of the assemblies. The final analysis was based on 548 species. To account for species with more than one remaining genome assembly, we calculated the average genome size for each species and plotted the average genome size grouped by superfamilies using R v 4.3.0^44^.

## Data Records

Raw sequence data, genome assemblies, and sample information (BioSample: SAMN38512512) are available on NCBI under Bioproject PRJNA1047044. All supporting data and materials are available as supplementary material. The removed contigs, helitron-related and ncRNA sequences, BUSCO gene coordinates, k-mer count histogram, and structural and functional annotations are available on Figshare (https://doi.org/10.6084/m9.figshare.24747036.v2).

## Technical Validation

### Quality of sequence and genome assembly

The raw sequence data included 5,194,461 HiFi ccs reads with an average read length of 13,685 bps (total length ca. 71 Gb) (**Supplementary Data 2**). Given the final assembly size of 2,586 Mb, we obtained a coverage of roughly 26.9X sequence depth. The k-mer based estimation reported by GenomeScope2 showed a slightly smaller genome size of 2,342 Mb with a sequence depth of 26X (**Supplementary Figure S3**). The estimate from backmap suggested a pick coverage at 25X with a 2,610 Mb (reported 2.61 Gb) genome, which was much closer to the final assembly size. The Hifiasm raw genome assembly included 2,960 Mb in 580 contigs (L50=37 and N50=22.14 Mb), with a high BUSCO completeness (97.2% complete, 95.1% single copy and 2.1% duplication) based on the endopterygota_odb10 database. After purging putatively duplicated haplotigs, only 184 contigs were kept and the assembly size was reduced to 2,586 Mb (L50=29 and N50=28.1 Mb). BUSCO completeness after purging remained intact, and only the duplication rate was reduced (97.3% complete, 96.3% single copy and 1.0% duplication), indicating that duplicated haplotigs were correctly identified and trimmed. During putative foreign DNA detection, we found that four contigs had blast hits from non-arthropod sequences (two from Ascomycota, one from Chordata, and one from Proteobacteria) (**Supplementary Figure S2**). Three of these were removed, but the contig that was blasted to Chordata (ptg000003l) was retained in the assembly because all 13 BUSCO genes on this contig were blasted to other lepidopteran species and the mean coverage and its GC content is similar to other contigs (**Supplementary Figure S2, Supplementary Data 1**). The final assembly thus consisted of 181 contigs and a total length of 2,586 Mb (L50=29 and N50=28.1 Mb), with high BUSCO completeness and low duplication rate (C:97.1% complete, 96.1% single copy and 1.0% duplication) (**Table 1**).

**Table 1.** Summary statistics and key characteristics of the *Druceiella hillmani* genome. Percentage of complete BUSCOs is given based on BUSCO v5.3.0 using the endopterygota_odb10 dataset.

| Assembly statistics |  |
| --- | --- |
| Genome size (bp) | 2,586,138,562 |
| Total contigs (n) | 181 |
| N50 contig length (bp) | 28,103,563 |
| L50 contigs (n) | 29 |
| GC content (%) | 0.3003 |
| Average sequence read coverage | 25x |
| **BUSCO scores** | C:97.1%[S:96.1%,D:1.0%],F:0.8%,M:2.1%,n:2124 |
| Augustus gene prediction models and functional annotation |  |
| Total predicted genes | 61,160 |
| Complete models | 59,371 |
| Monoexonic genes | 13,595 |
| Multiexonic genes | 47,527 |
| Functionally annotated genes (eggNOG) | 26,790 |
| Functionally annotated genes (blast) | 26,865 |
| **BUSCO scores** | C:89.4%[S:85.5%,D:3.9%],F:3.5%,M:7.1%,n:2124 |
| Repeat and transposable elements |  |
| Total repeat content (%) | 0.3759 |
| LINEs | 0.0457 |
| LTR elements | 0.0178 |
| DNA elements | 0.0538 |
| Unclassified repeats | 0.1768 |
| non-coding RNA (Infernal) | 0.0001 |
| helitron-related sequences | 0.0088 |
| BioProject Accession | PRJNA1047044 |
| BioSample Accession | SAMN38512512 |

### Genome sizes of Lepidoptera

We compared the genome size of *Druceiella hillmani* with that of 548 other lepidopteran species from 45 families and 21 superfamilies (**Figure 2, Supplementary Data 3**). The average genome size among superfamilies ranged from 300 Mb (Choreutoidea) to 2,586 Mb (Hepialoidea, this study), but the majority of the included species had a genome size well below 1,000 Mb. Before this study, genome sizes had been characterized for four hepialid species (*Hepialus humuli* = 2,610 Mb, *Triodia sylvina* = 1,800-3,080 Mb, *Thitarodes* sp. = 3,166 Mb, and *Phymatopus californicus* = 1,580 Mb) using short-read *de novo* assembly and cytogenetics methods^5,6,45^. With currently known genome sizes ranging from 1,580 Mb to 3,166 Mb, hepialids include the largest lepidopteran genomes known so far. This suggests that there may have been an ancestral large-scale genome expansion within the family. Genome size expansions are less common than genome contractions and their evolutionary implications are often not clear^46–49^, even though they have been suggested to influence host range, reproductive fitness, and morphological traits in different taxa^46,50–52^. Recently, genome size expansions in caddisflies (Trichoptera), the sister taxon of Lepidoptera, were related to more diverse and unstable habitats^53^. Interestingly, most ghost moth caterpillars live at high elevations and have a distinct subterranean lifestyle and share several ecological characteristics with the butterfly *Parnassius apollo*, which has the largest known genome among butterflies and is one of very few butterflies that pupates in the soil in high-elevation areas^8^. It is worth further investigating whether these exceptional ecological features played a role in the genome expansions in both lineages. Ancestral genome expansions have also been suggested to increase diversification^47,53^, and the increased genome size in Hepialidae might have led to increased diversity in this family compared to closely related families.

**Figure 2.**
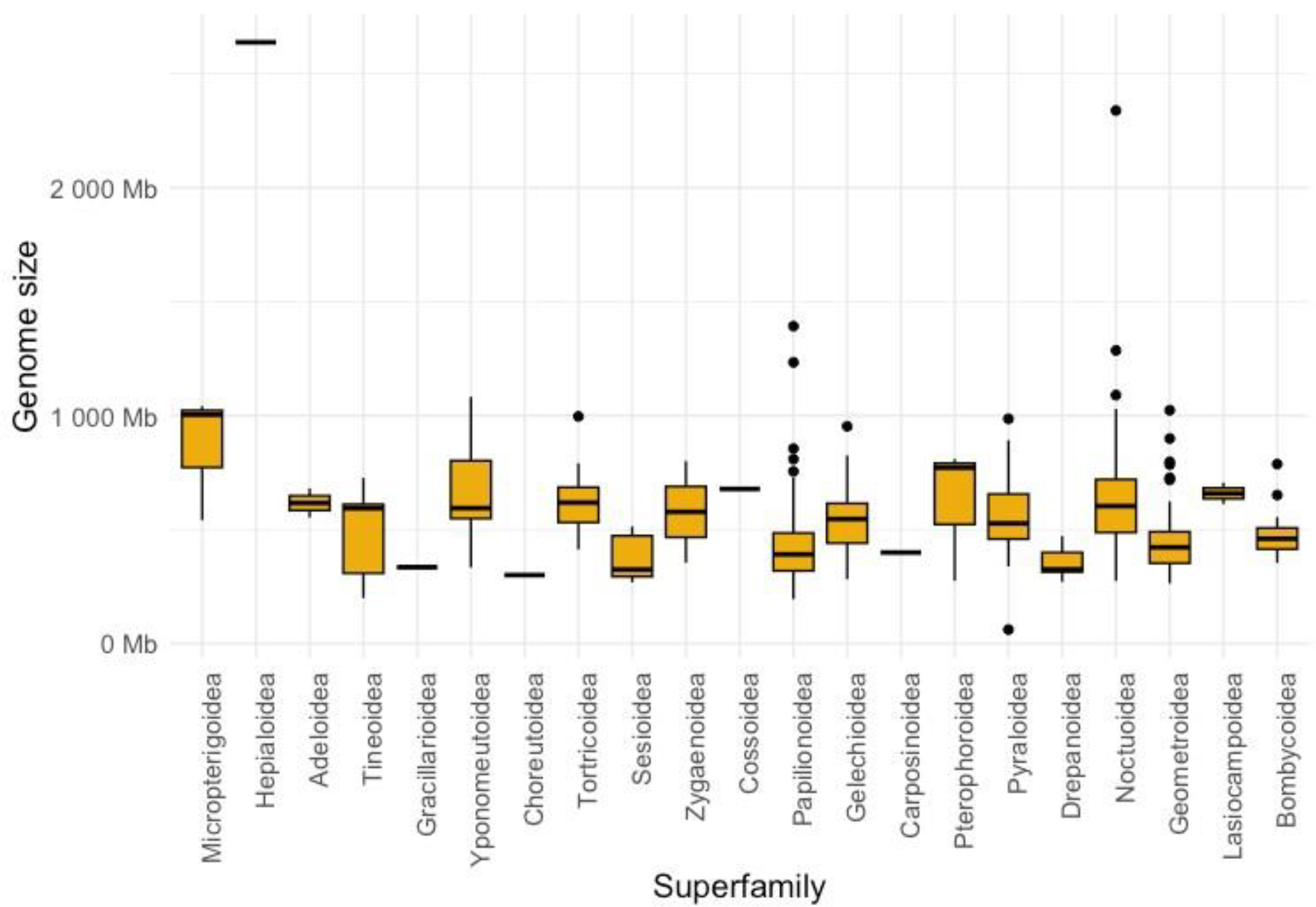
Comparison of genome sizes in 548 species of Lepidoptera. Genome Assemblies for this analysis were chosen based on a BUSCO completeness >80% and duplication rate <5%. Box plots represent the mean genome size across 21 superfamilies within Lepidoptera. The genome of *Druceiella hillmani* was among the largest of the 548 available high-quality Lepidoptera genomes that were compared.

There are multiple mechanisms potentially underlying the large genome assembly, both evolutionary and technical in nature. First, evidence of genome duplications has been found in many insect lineages, including in Lepidoptera^54^. However, we consider it unlikely that the expanded genome of *D. hillmani* is the result of recent genome duplication. Indeed, the low BUSCO duplication rate and the clear two-peak k-mer histogram from the *D. hillmani* genome disprove the contribution of a recent genome duplication to the expanded genome in this species (**Supplementary Figure S3**). We cannot rule out the possibility of a more ancient genome duplication, earlier in the evolutionary history of Hepialidae, as genomic data from other hepialid species will be needed to test for ancient genome duplications. We also consider non-target DNA from other species, such as parasitoids or gut contents, an unlikely explanation for the large genome assembly, as most annotated genes were blasted to other lepidopteran species and none of our contigs are dominated by genes that blast to species outside of Lepidoptera (**Figure 3, Supplementary Data 4**).

**Figure 3.**
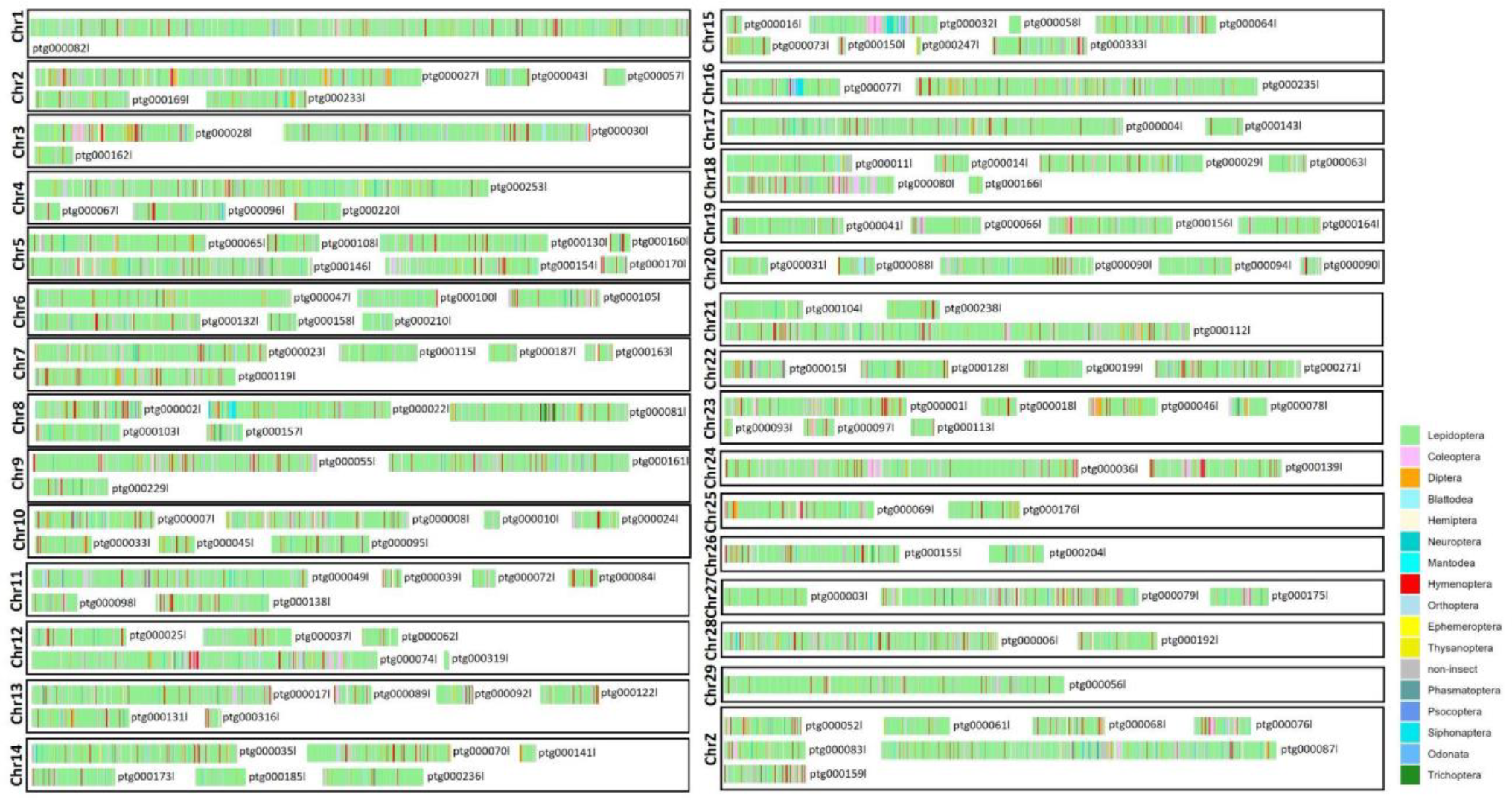
Gene map of coding gene blast results against the RefSeq non-redundant (nr) protein database from NCBI. The best hit (defined by lowest e-value) was selected for each gene, and the order of the corresponding species was mapped to the contigs. Most genes had their best blast hit to lepidopteran species, and no contig was found to harbor genes dominantly from a different order than Lepidoptera.

### Structural and functional genome annotation

The final assembly was soft-masked with repeat sequences based on the repeat library from RepeatModeler2 and Repbase Arthropoda database. Repeat elements accounted for 37.59% of the genome (943.85 Mbp), including 4.57% LINEs, 1.78% LTR elements, 5.38% DNA elements, and 17.68% unclassified elements (**Supplementary Table S1**). In addition, we found 2,275 Helitron-related sequences contributing to 0.88% (22,683,504 bp) of the genome and 1,990 non-coding RNA (ncRNA) genes accounting for 0.012% (312,251 bp) of the genome. Repetitive elements (REs) have repeatedly been shown to be an important driver of genome expansions^46,53,55,56^, but surprisingly contribute to a lower proportion of the *D. hillmani* genome (37.59%) than expected. This unexpectedly low proportion of REs could be caused by the inability of current RE detection software to detect ancient REs where mutations have accumulated, as has been suggested before55,57,58.

Protein coding gene models predicted from the BRAKER3 pipeline using GeneMark-EP+ module resulted in large gene sets, where the braker model predicts 61,589 transcripts from 59,148 genes, and the Augustus model predicts 61,122 transcripts from 58,936 genes. The larger number of genes predicted in the *Druceiella hillmani* genome assembly, compared to other lepidopteran genomes, partially reflects the large genome size. The Augustus gene model has BUSCO scores of 89.4% complete, 85.5% single copy and 3.9% duplication. The lower BUSCO completeness is likely the result of the lack of transcriptomic data in gene model training. Among the 61,122 Augustus transcripts, 13,595 are monoexonic and 47,527 are multiexonic, and the average overall gene size is 7,667 bp (**Supplementary Table S2**). The functional annotation resulted in 26,790 annotated genes with EggNOG function, 12,679 with gene ontology term (GO term), and 26,865 genes that are blasted to sequences in NCBI nr database (**Table 1**). The new genome made available in this study allows for more detailed study of this genome size expansion and gene evolution.

## Code availability

All bioinformatic tools for data analysis were used according to their manuals, and are cited in the relevant parts of the methods section. No custom code was used.

## Acknowledgements

This project was financially supported in part by the National Science Foundation (NSF) grant number EF-2217159 to AYK. Paul Frandsen (Brigham Young University) and Andrew Mongue (University of Florida) provided useful discussions. We thank Taylor Pierson and Christian Couch (UF MGCL) for helping with sample collection and preparation. Jesse Barber and Pamela Rivera (Boise State University) helped with collecting and transportation permits. The BYU sequencing core and UF’s HiPerGator cluster provided sequencing resources and computational assistance, respectively.

## Author contributions

YMW: Project design, sample preparation, data analysis, manuscript writing

AIC Figure preparation, manuscript writing

BF: Manuscript writing and editing

JIM: Identification, supplementary material preparation, manuscript writing

DP: Manuscript writing and editing

AS: Project design, manuscript writing and editing

AYK: Sample collection, funding, manuscript writing

## Supplementary information

Supporting information: Supplementary Tables and Supplementary Figures Supplementary Data 1: Assembly contig summary

Supplementary Data 2: HiFi sequence report Supplementary Data 3: Lepidoptera genome size data Supplementary Data 4: Summary of gene blast result

