## supporting_information for "A high-quality genome assembly of the ghost moth *Druceiella hillmani* provides new evidence of genome size augmentation in Hepialidae"

**Supplementary Table S1.** Summary of repeat elements in the *Druceiella hillmani* genome from RepeatMasker

| Repeat elements | Family | Number of elements | Length occupied (bp) | Percentage of sequence |
|---|---|---|---|---|
| SINEs | | 0 | 0 | 0.00% |
| | ALUs | 0 | 0 | 0.00% |
| | MIRs | 0 | 0 | 0.00% |
| LINEs | | 389,454 | 118,071,727 | 4.57% |
| | LINE1 | 0 | 0 | 0.00% |
| | LINE2 | 79,256 | 23,837,622 | 0.92% |
| | L3/CR1 | 76,747 | 24,496,779 | 0.95% |
| LTR elements | | 160,513 | 45,974,112 | 1.78% |
| | ERVL | 0 | 0 | 0.00% |
| | ERVL-MaLRs | 0 | 0 | 0.00% |
| | ERV_classI | 0 | 0 | 0.00% |
| | ERV_classII | 0 | 0 | 0.00% |
| DNA elements | | 421,680 | 139,006,240 | 5.38% |
| | hAT-Charlie | 5,337 | 2,685,584 | 0.10% |
| | TcMar-Tigger | 3,713 | 1,437,454 | 0.06% |
| Unclassified | | 2,277,069 | 457,102,022 | 17.68% |
| Total interspersed repeats | | | 760,154,101 | 29.39% |
| Small RNA | | 6,858 | 1,454,523 | 0.06% |
| Satellites | | 0 | 0 | 0.00% |
| Simple repeats | | 772,641 | 47,649,454 | 1.84% |
| Low complexity | | 115,212 | 5,573,392 | 0.22% |

**Supplementary Table S2.** Summary of the gene model from the Augustus prediction in BRAKER3 pipeline

| **Description** | **Statistics** |
| --- | --- |
| Number of genes | 61,160 |
| Number of monoexonic genes | 13,595 |
| Number of multiexonic genes | 47,527 |
| Number of positive strand genes | 30,085 |
| Number of positive strand monoexonic genes | 6,730 |
| Number of positive strand multiexonic genes | 23,317 |
| Number of negative strand genes | 31,075 |
| Number of negative strand monoexonic genes | 6,865 |
| Number of negative strand multiexonic genes | 24,210 |
| Average overall gene size | 7,667.15 |
| Median overall gene size | 3,187 |
| Average overall CDS size | 753.413 |
| Median overall CDS size | 435 |
| Average overall exon size | 218.838 |
| Median overall exon size | 165 |
| Average size of monoexonic genes | 497.332 |
| Median size of monoexonic genes | 318 |
| Largest monoexonic gene | 12,906 |
| Smallest monoexonic gene | 110 |
| Average size of multiexonic genes | 9,723.85 |
| Median size of multiexonic genes | 5407 |
| Largest multiexonic gene | 197,036 |
| Smallest multiexonic gene | 240 |
| Average size of multiexonic CDS | 827.267 |
| Median size of multiexonic CDS | 480 |
| Largest multiexonic CDS | 81,981 |
| Smallest multiexonic CDS | 21 |
| Average size of multiexonic exons | 199.616 |
| Median size of multiexonic exons | 159 |
| Average size of multiexonic introns | 2,829.23 |
| Median size of multiexonic introns | 1641 |
| Average number of exons per multiexonic gene | 4.144 |
| Median number of exons per multiexonic gene | 3 |
| Largest multiexonic exon | 14,886 |
| Smallest multiexonic exon | 3 |
| Most exons in one gene | 294 |
| Average number of introns per multiexonic gene | 3.144 |
| Median number of introns per multiexonic gene | 2 |
| Largest intron | 53,398 |
| Smallest intron | 16 |
| Number of complete models | 59,371 |
| Number of 5' only incomplete models | 997 |
| Number of 3' only incomplete models | 676 |
| Number of 5' and 3' incomplete models | 78 |

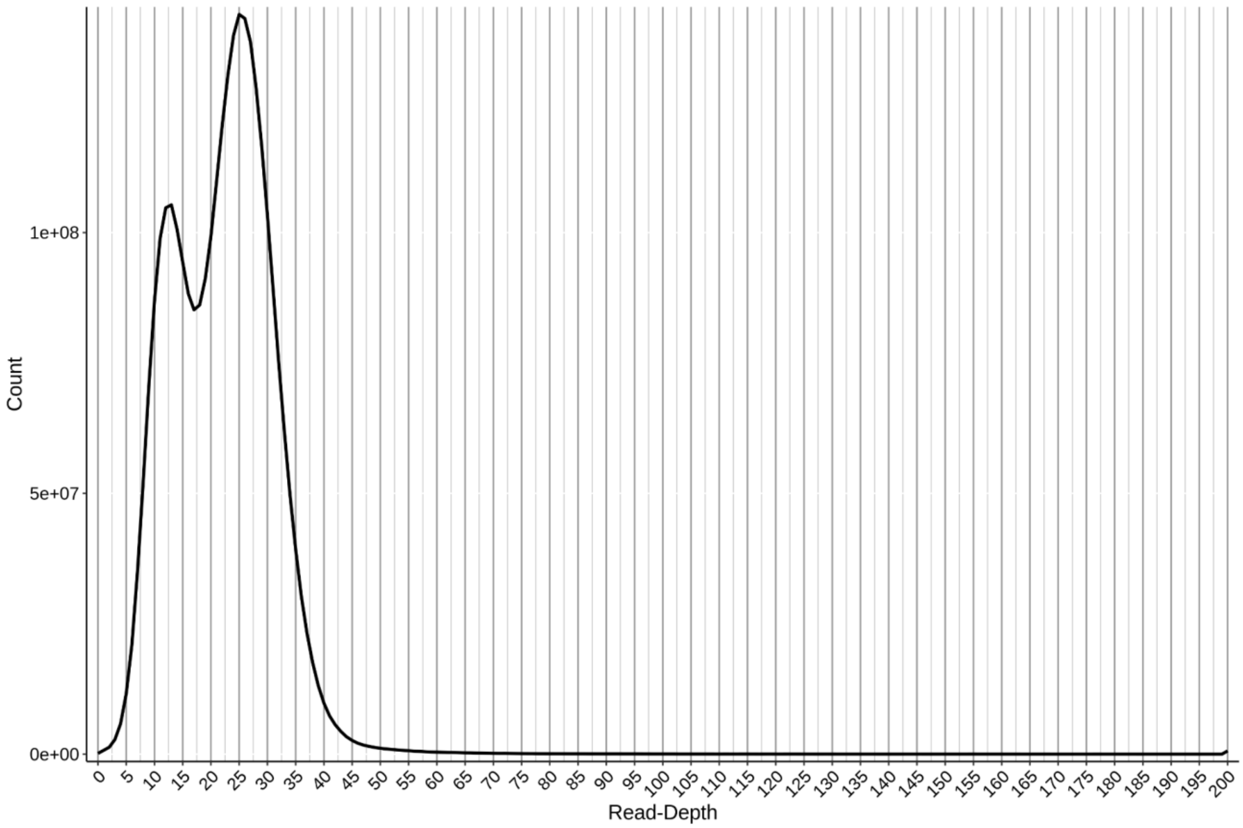


**Supplementary Figure S1.** Mapped coverage histogram generated from the purge_haplotigs pipeline. The read depth was calculated based on the mapping results of minimap2 where the raw sequence reads were mapped to the raw Hifiasm assembly.

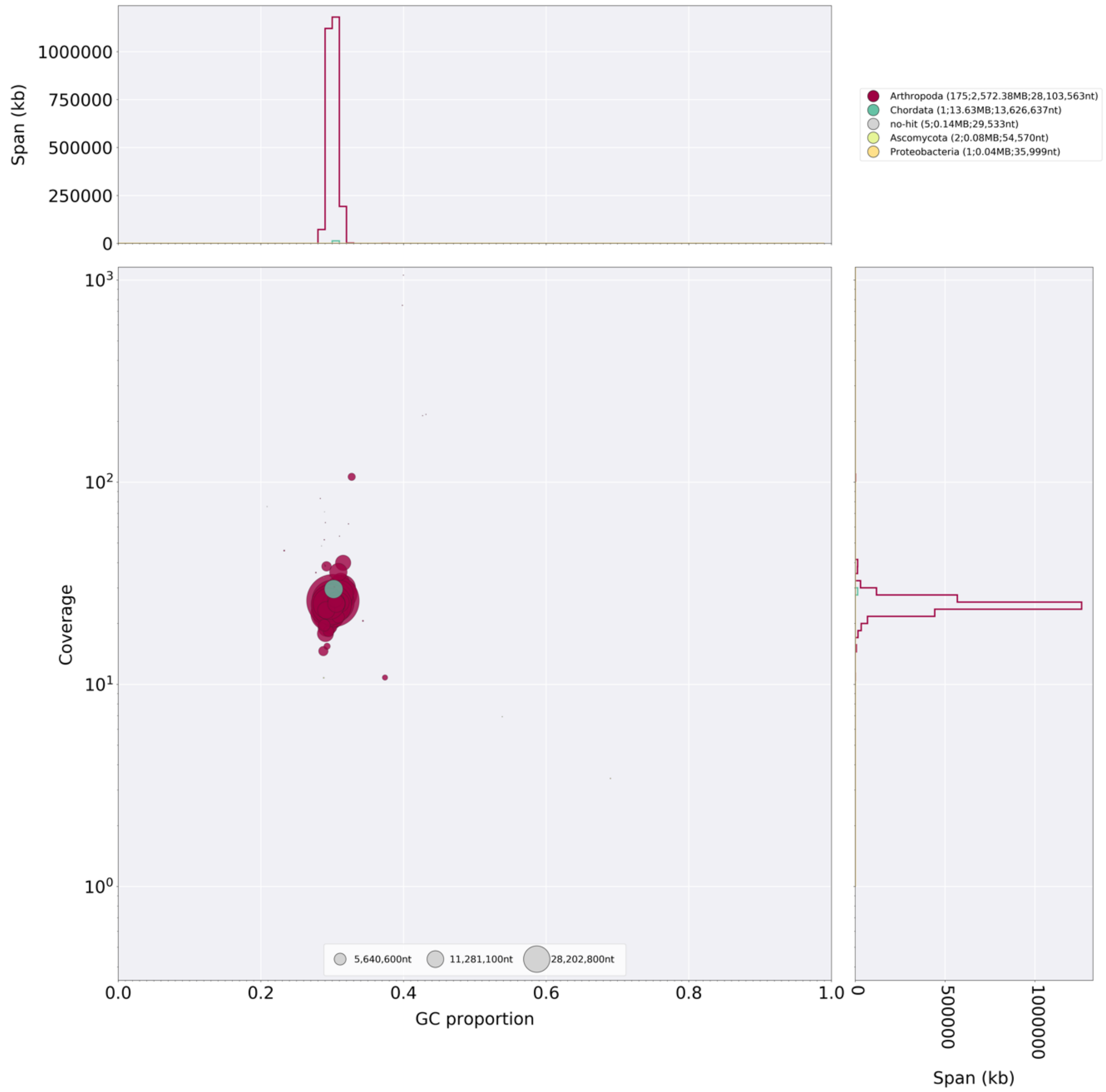


**Supplementary Figure S2.** BlobPlot of the *Druceiella hillmani* purged genome assembly. Red dots show contigs with best blast hits to Arthropoda; light green to Ascomycota; dark green dot Chordata; yellow dot to Proteobacteria; and gray dots had no hits. Contigs with best hit to Ascomycota and Proteobacteria were removed and the contig with best hit to Chordata was kept.

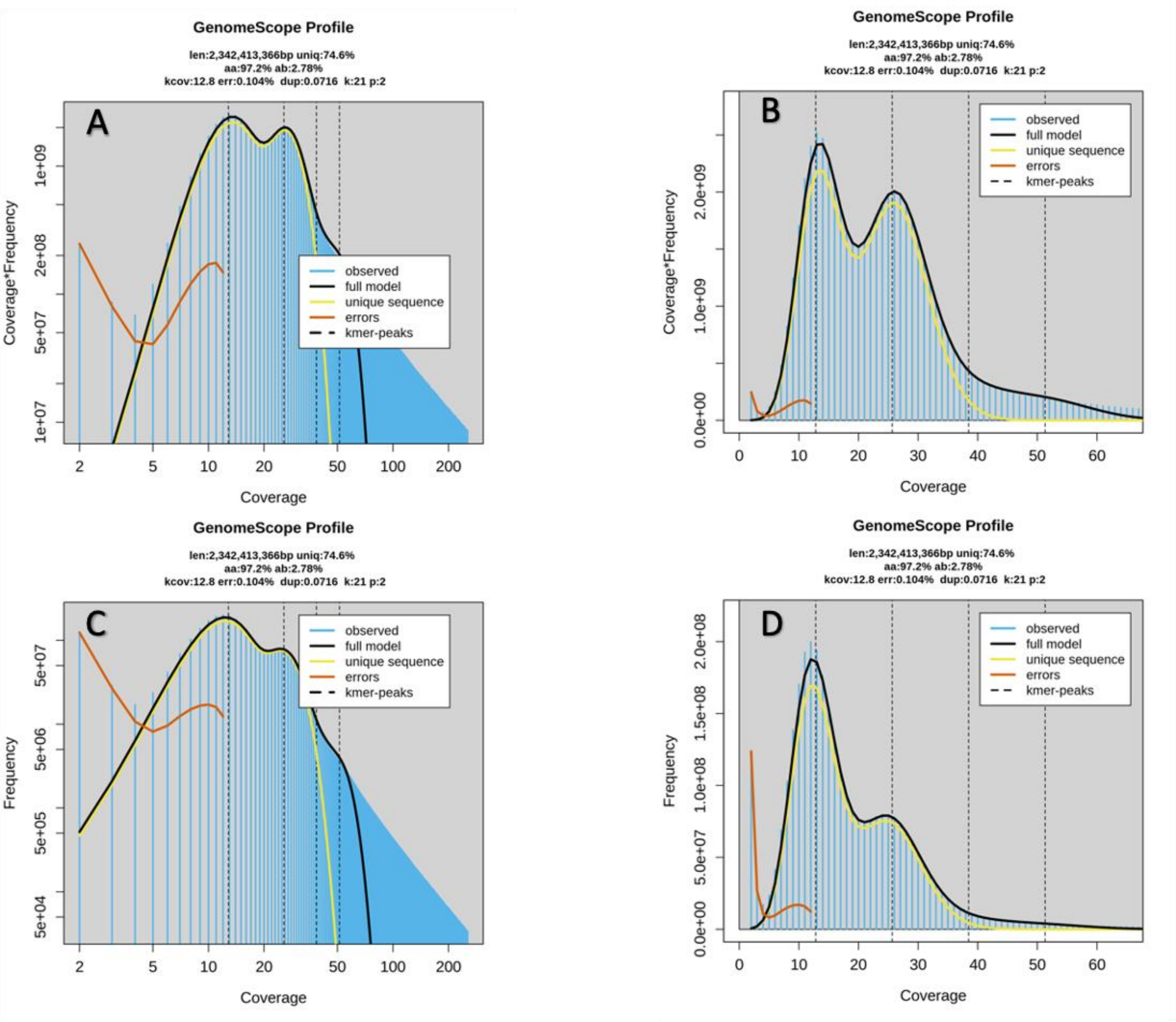


**Supplementary Figure S3.** K-mer distribution calculated using KMC with k-mer size of 21 bps. The fitting model (black line) is created using GenomeScope2. The estimated genome size is around 2,342 Mb with k-mer coverage being 12.8X (homozygous peak coverage).
