## supplementary_data2 for "A high-quality genome assembly of the ghost moth *Druceiella hillmani* provides new evidence of genome size augmentation in Hepialidae"

### Report for dataset Lep 89533-Cell2 (all samples)

### Dataset 6b28562e-425d-4469-8bab-271d2e23fb19

##### Summary

| | |
|---|---|
| **Name** | Lep 89533-Cell2 (all samples) |
| **Created At** | 2023-07-16 14:20:43.187 |
| **Number of Records** | 4995450 |
| **Total Length** | 68367713586 |
| **Movie ID** | m84100_230715_010611_s2 |
| **ICS Version** | 12.0.0.179648 |
| **Well Sample** | Lep 89533 |
| **Biological Sample** | Lep 89533 |
| **Barcode Name** | default |

### CCS Processing

##### Summary

| | |
|---|---|
| **ZMWs input** | 20,097,756 |
| **ZMWs pass filters** | 5,672,860 |
| **ZMWs fail filters** | 14,424,896 |
| **ZMWs shortcut filters** | 0 |
| **ZMWs with tandem repeats** | 98,199 |
| **Below SNR threshold** | 202,210 |
| **Median length filter** | 0 |
| **Lacking full passes** | 13,593,403 |
| **Heteroduplex insertions** | 149,005 |
| **Coverage drops** | 5,483 |
| **Insufficient draft cov** | 100,535 |
| **Draft too different** | 1 |
| **Draft generation error** | 319,437 |
| **Draft above --max-length** | 0 |
| **Draft below --min-length** | 167 |
| **Reads failed polishing** | 18,816 |
| **Empty coverage windows** | 20,693 |
| **CCS did not converge** | 9,954 |
| **CCS adapter concatenation** | 1 |
| **CCS adapter palindrome** | 81 |
| **CCS adapter residue** | 141 |
| **CCS below minimum RQ** | 4,969 |
| **Unknown error** | 0 |
| **ZMWs missing adapters** | 387,377 |

### Adapter Report

#### Summary

| | |
|---|---|
| **Adapter Dimers (0-10bp) %** | 0 |
| **Short Inserts (11-100bp) %** | 0 |
| **Local Base Rate** | 1.79 |

### CCS Analysis Report

#### Summary

| | |
|---|---|
| **HiFi Reads** | 5,194,461 |
| **HiFi Yield (bp)** | 71,087,217,156 |
| **HiFi Read Length (mean, bp)** | 13,685 |
| **HiFi Read Length (median, bp)** | 13,037 |
| **HiFi Read Length N50 (bp)** | 13,619 |
| **HiFi Read Quality (median)** | Q31 |
| **HiFi Read Quality (median)** | 31 |
| **HiFi Number of Passes (mean)** | 8 |

#### HiFi Read Length Summary

| Read Length (bp) | Reads | Reads (%) | Yield (bp) | Yield (%) |
|---|---|---|---|---|
| 0 | 5194461 | 100 | 71087217156 | 100 |
| 5,000 | 5190332 | 100 | 71076141522 | 100 |
| 10,000 | 5149082 | 99 | 70696834176 | 99 |
| 15,000 | 1392291 | 27 | 24350856353 | 34 |
| 20,000 | 183627 | 4 | 3948325916 | 6 |
| 25,000 | 2845 | 0 | 74239402 | 0 |
| 30,000 | 93 | 0 | 3100495 | 0 |
| 35,000 | 23 | 0 | 889742 | 0 |
| 40,000 | 5 | 0 | 221033 | 0 |

#### HiFi Read Quality Summary

| Read Quality (Phred) | Reads | Reads (%) | Yield (bp) | Yield (%) |
|---|---|---|---|---|
| Q20 | 5194461 | 100 | 71087217156 | 100 |
| Q30 | 2772323 | 53 | 36882379708 | 52 |
| Q40 | 672353 | 13 | 8281887109 | 12 |
| Q50 | 59480 | 1 | 666842046 | 1 |

**HiFi Read Length Distribution**

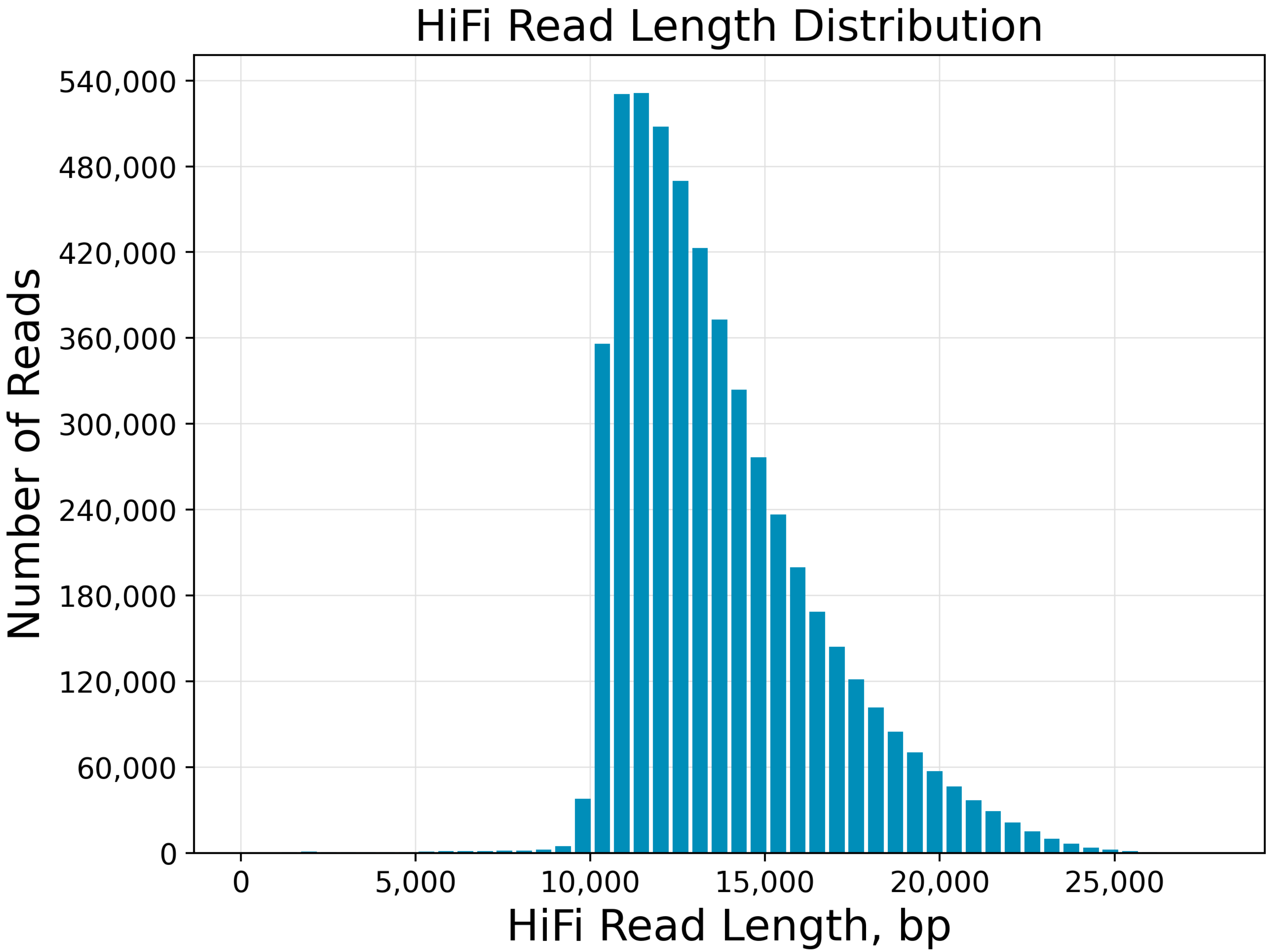

**Yield by HiFi Read Length**

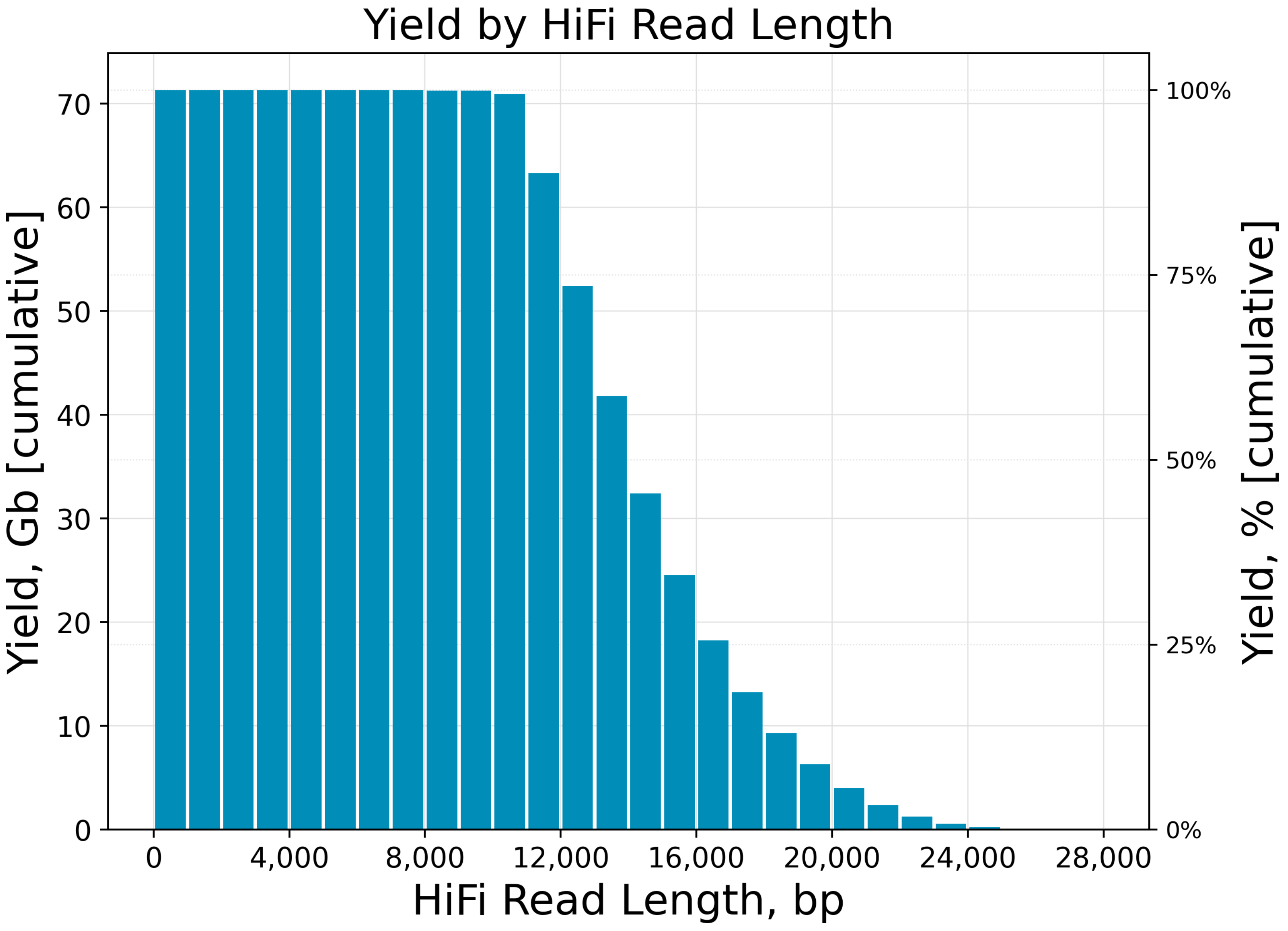

**Read Length Distribution**

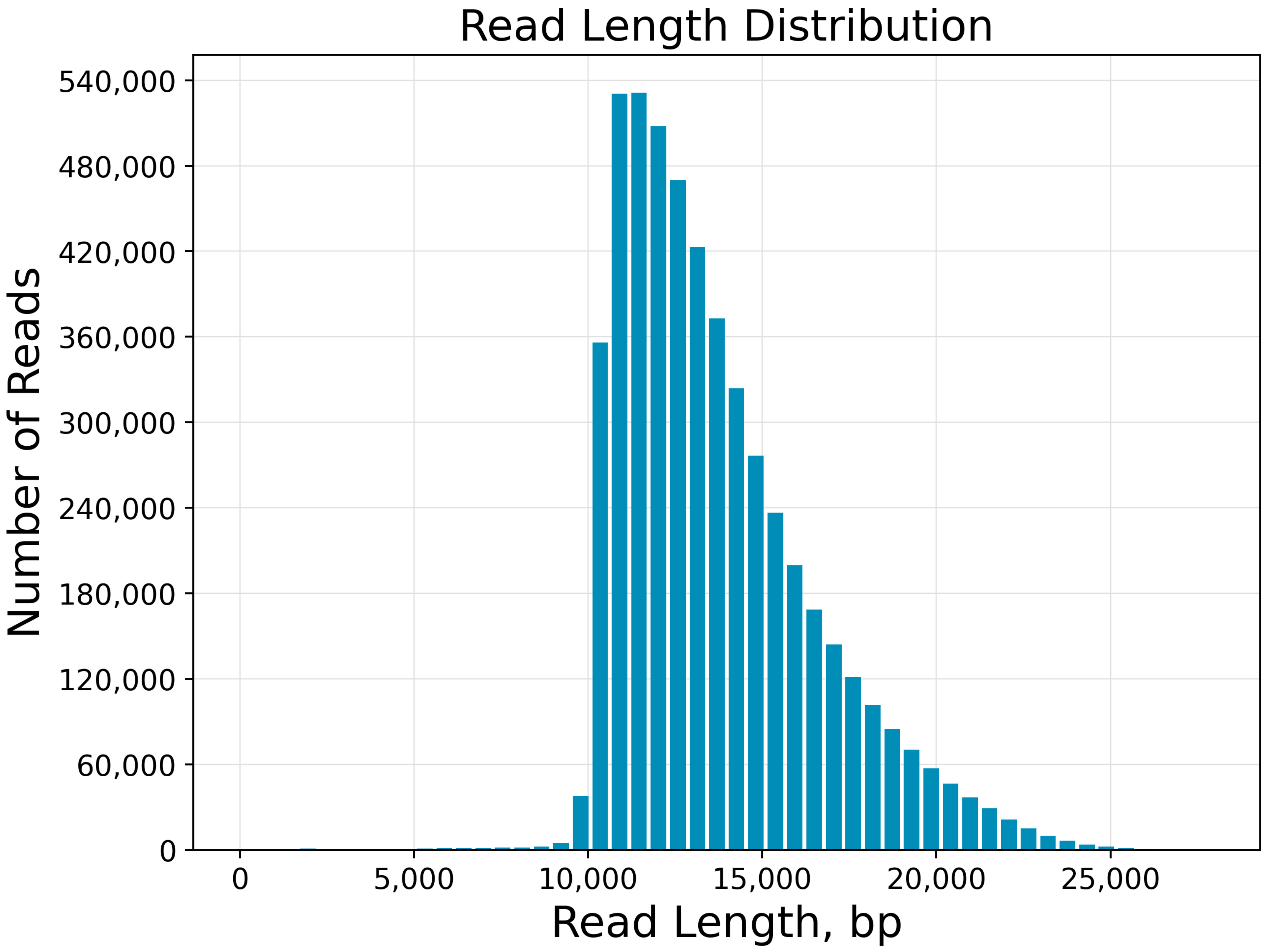

**Number of Passes**

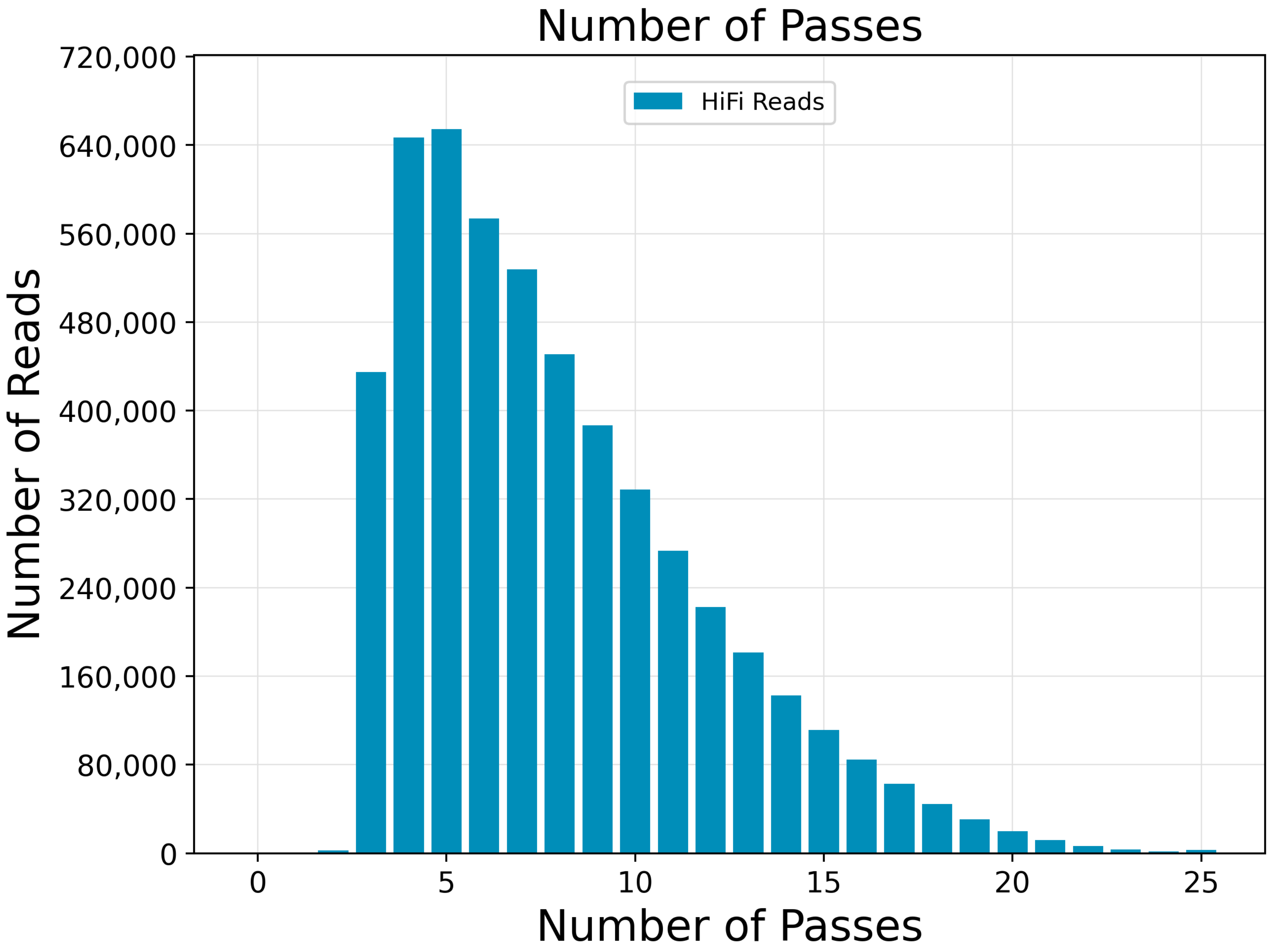

**Read Quality Distribution**

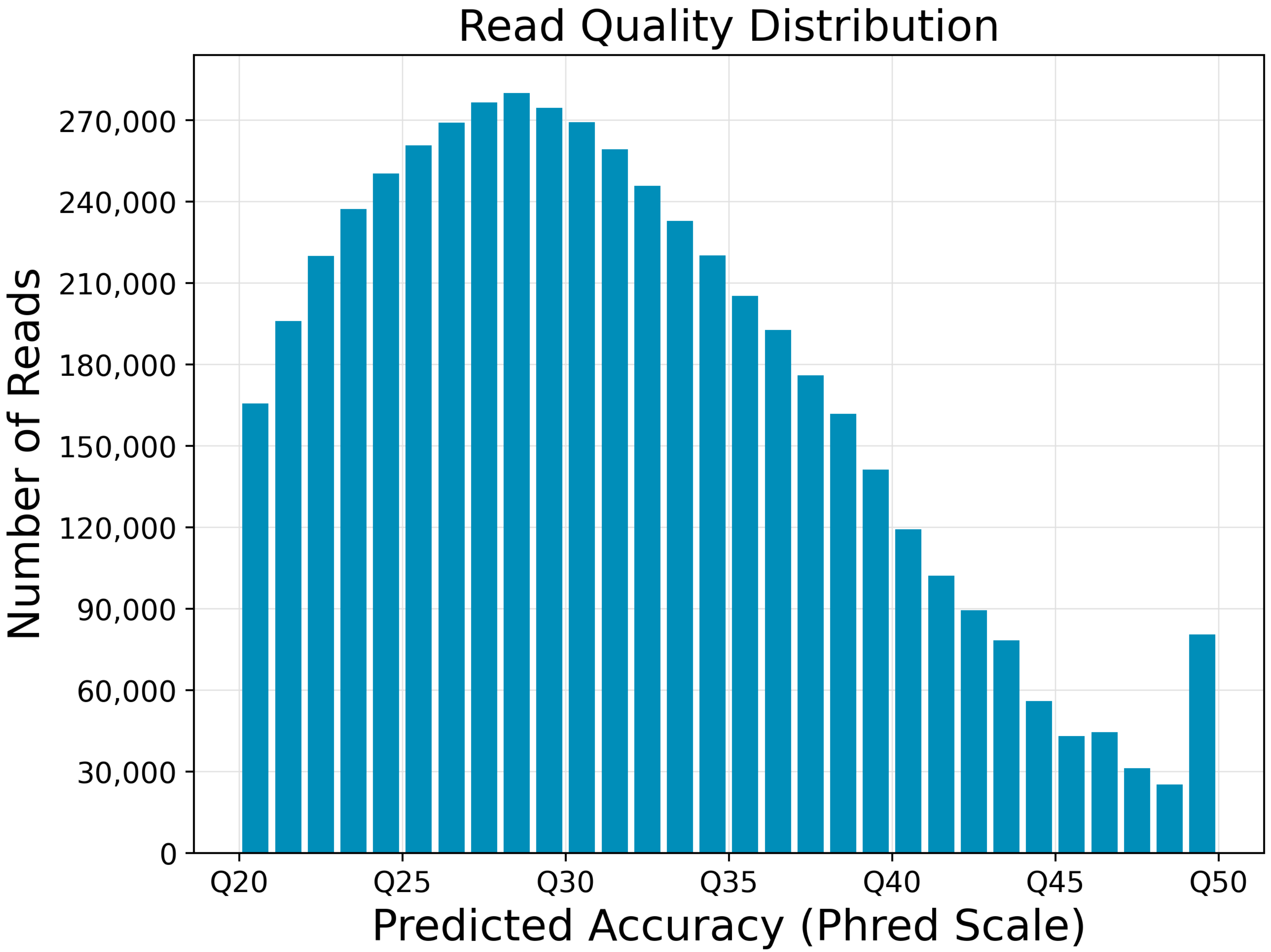

**Predicted Accuracy vs. Read Length**

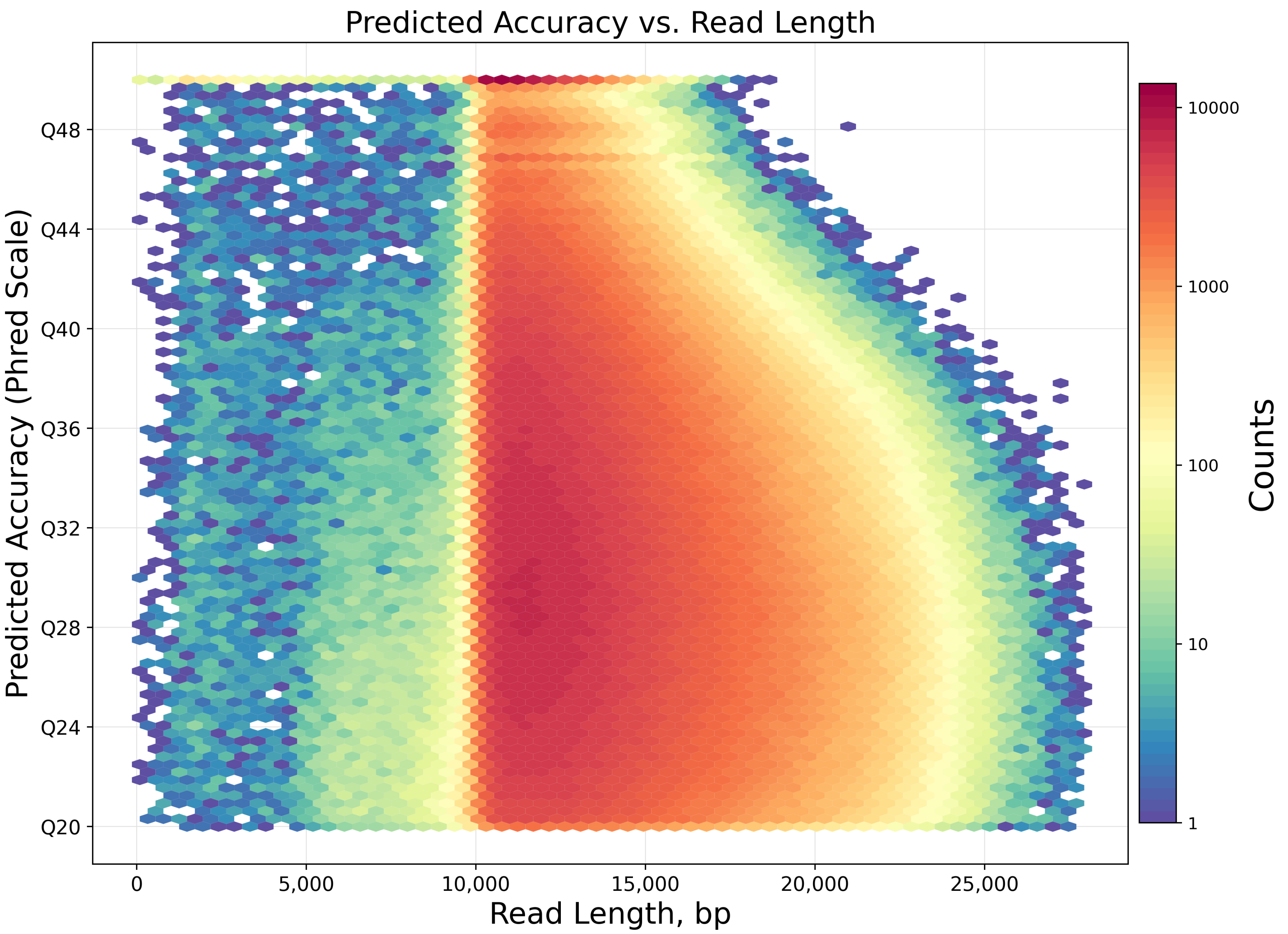

### Control Report

#### Summary

| | |
|---|---|
| **Number of Control Reads** | 660 |
| **Control Read Length Mean** | 46,256 |
| **Control Read Concordance Mean** | 0.90 |
| **Control Read Concordance Mode** | 0.91 |

**Control Polymerase RL**

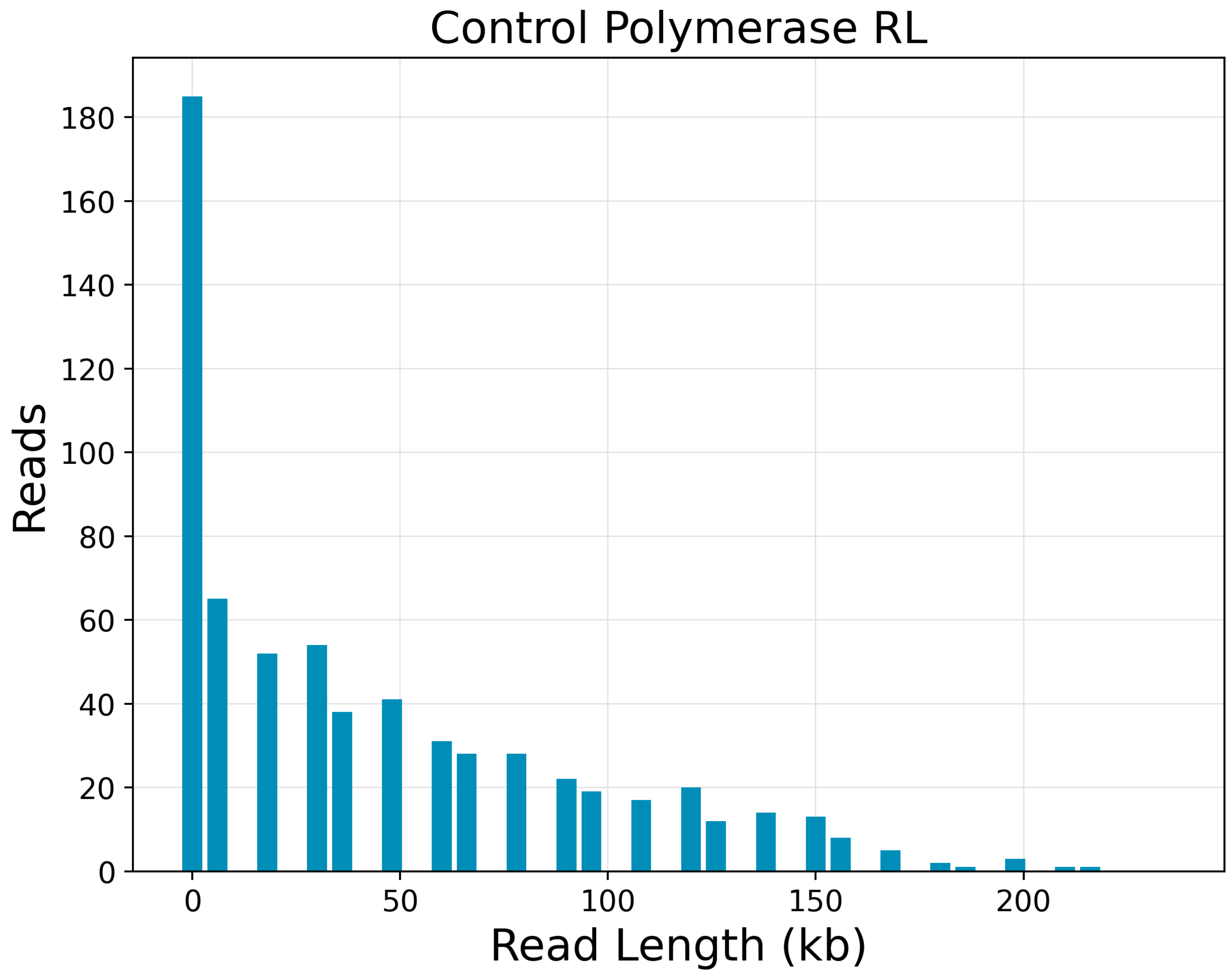

Control Polymerase RL
Reads
180
160
140
120
100
80
60
40
20
0
0
50
100
150
200
Read Length (kb)

#### Control Concordance

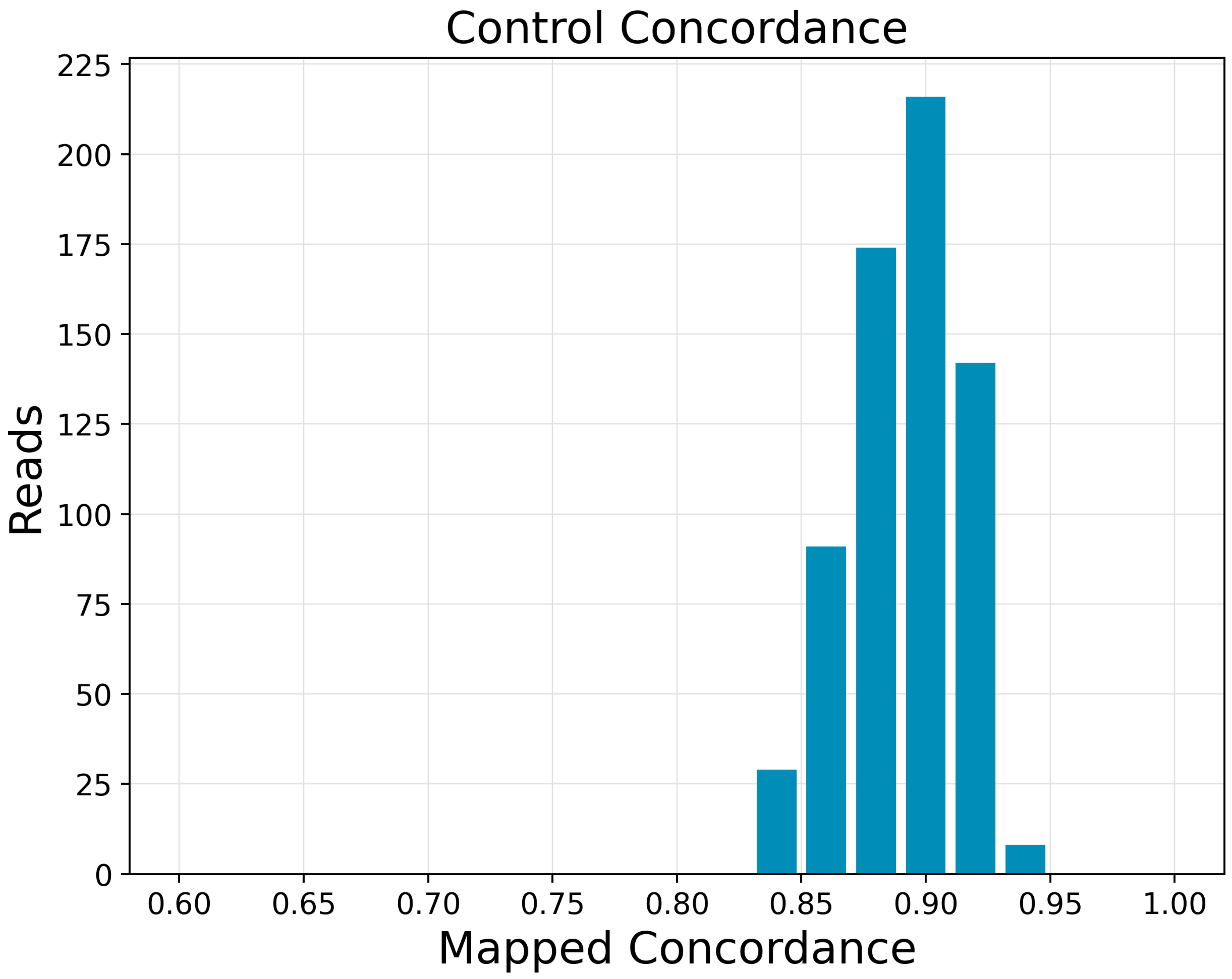
Control Concordance
Reads
Mapped Concordance
225
200
175
150
125
100
75
50
25
0
0.60
0.65
0.70
0.75
0.80
0.85
0.90
0.95
1.00

### 5mC CpG Report

#### CpG Methylation in Reads

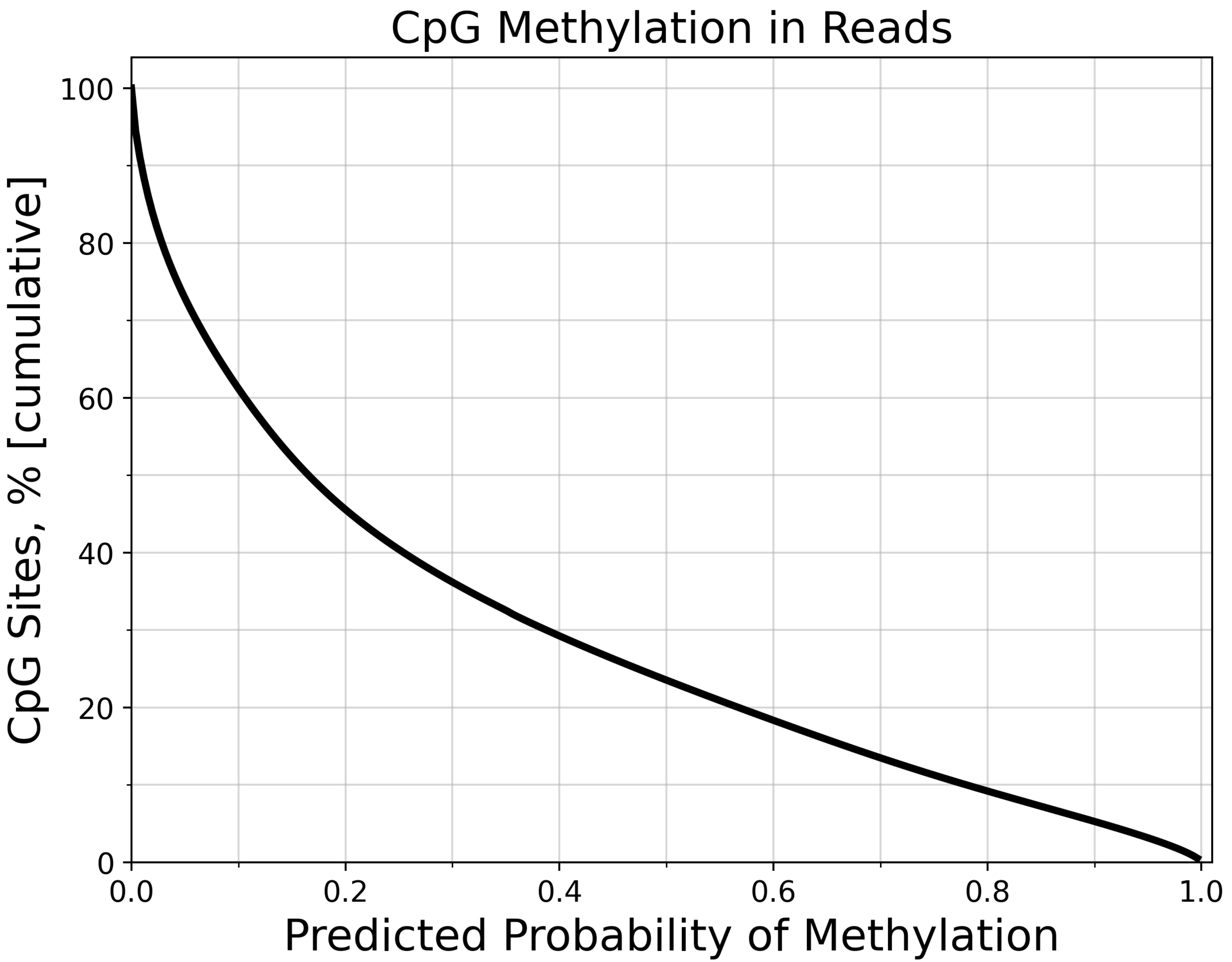

CpG Methylation in Reads
CpG Sites, % [cumulative]
100
80
60
40
20
0
0.0
0.2
0.4
0.6
0.8
1.0
Predicted Probability of Methylation

#### CpG Methylation in Reads (Histogram)

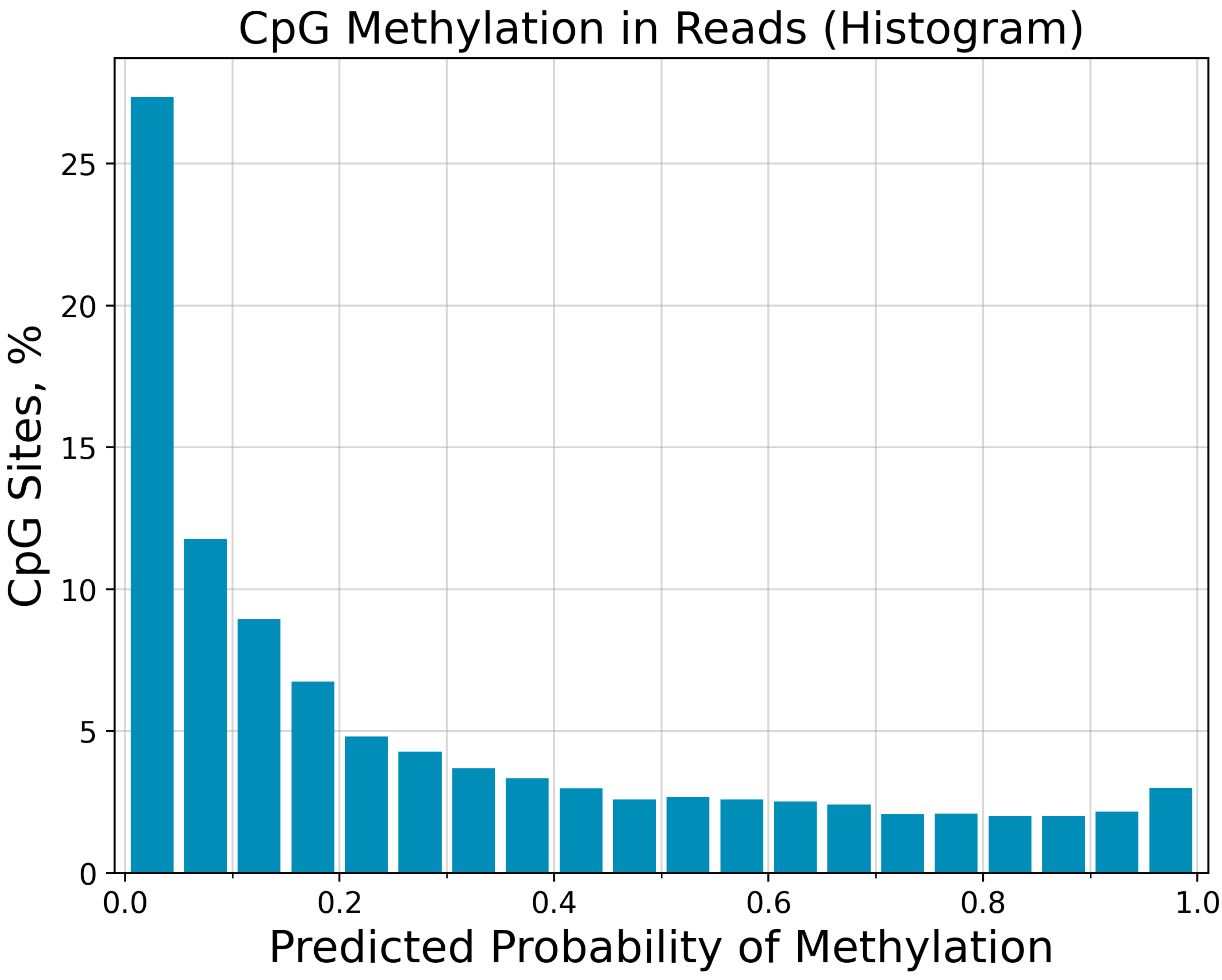

CpG Methylation in Reads (Histogram)
CpG Sites, %
Predicted Probability of Methylation
0
5
10
15
20
25
0.0
0.2
0.4
0.6
0.8
1.0

### Loading Report

#### Summary

| | |
|---|---|
| **Productive ZMWs** | 16,777,216 |
| **Productivity 0** | 3,927,778 |
| **Productivity 1** | 20,205,425 |
| **Productivity 2** | 1,032,621 |

#### Loading Statistics

| Collection Context | Productive ZMWs | Productivity 0 | (%) | Productivity 1 | (%) | Productivity 2 | (%) | Loading type |
|---|---|---|---|---|---|---|---|---|
| m84100_230715_010611_s2 | 16777216 | 3927778 | 15.61 | 20205425 | 80.29 | 1032621 | 4.10 | Workflow_Kestrel.py |

**HQ Region Filtering Evaluation**

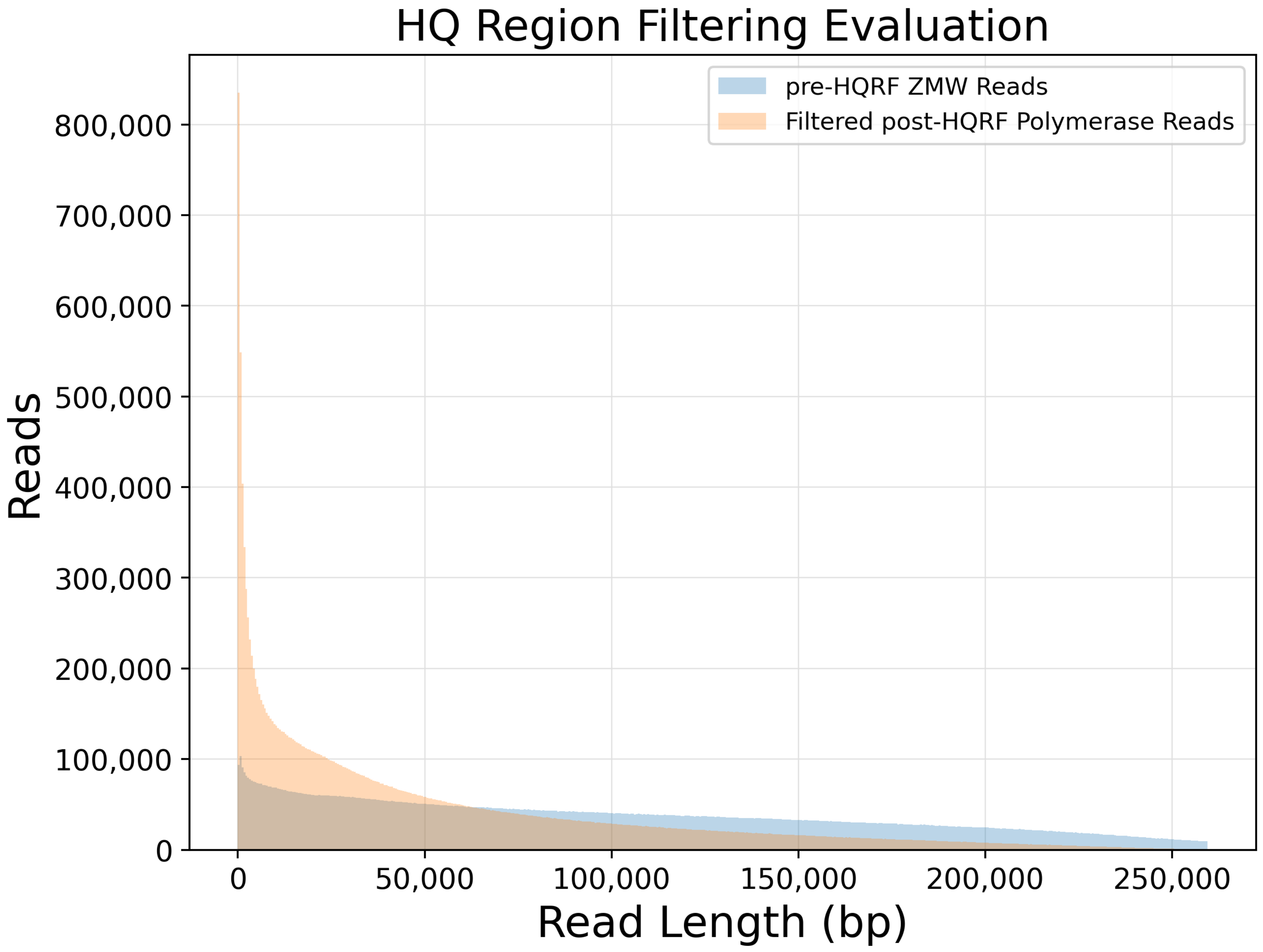

### Raw Data Report

#### Summary

| | |
|---|---|
| **Polymerase Read Bases** | 1,042,893,191,193 |
| **Polymerase Reads** | 20,204,765 |
| **Polymerase Read Length (mean)** | 51,616 |
| **Polymerase Read N50** | 105,250 |
| **Longest Subread Length (mean)** | 17,059 |
| **Longest Subread N50** | 22,250 |
| **Unique Molecular Yield** | 320,789,413,888 |

**Polymerase Read Length**

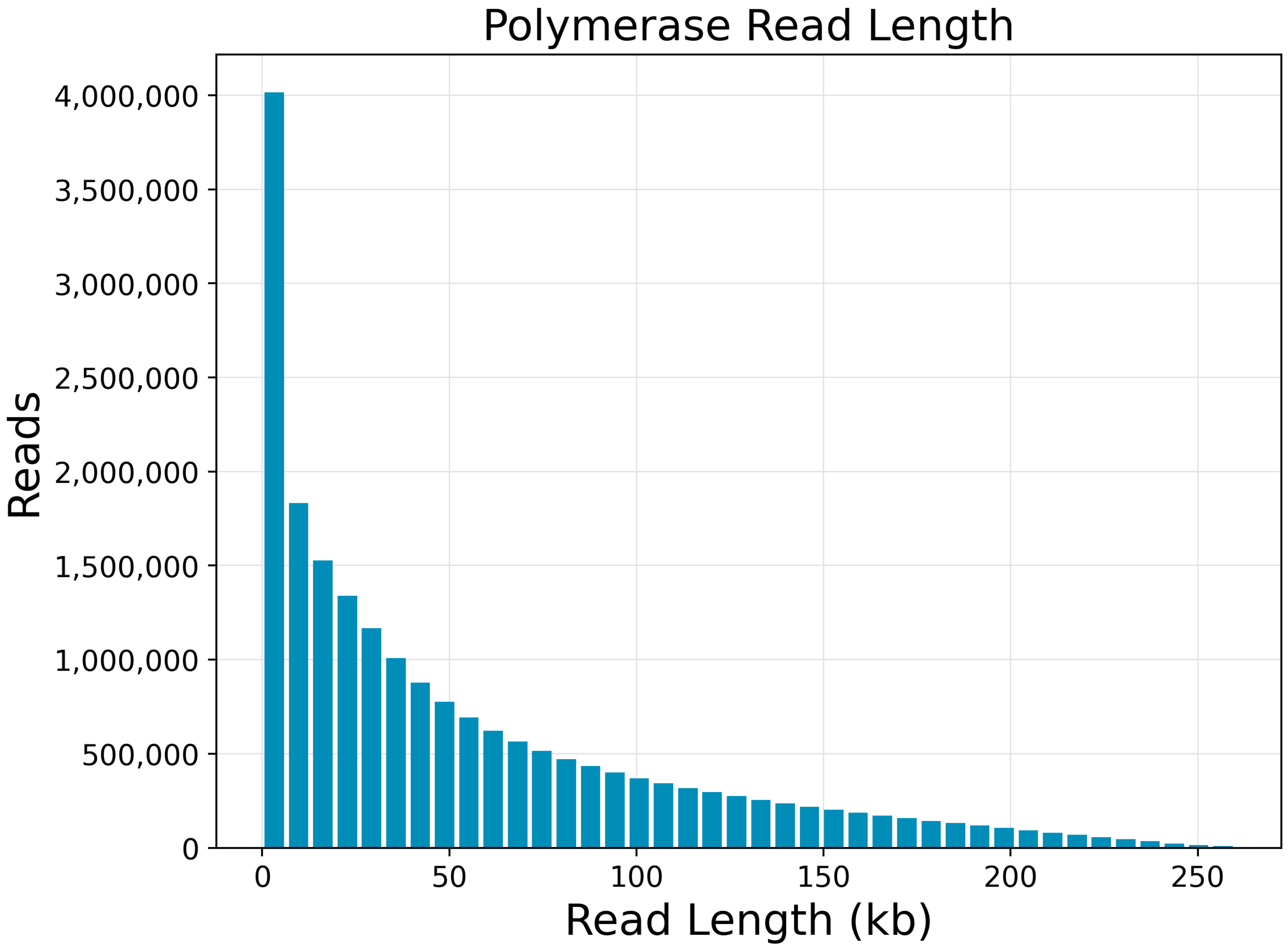

**Longest Subread Length Versus Polymerase Read Length**

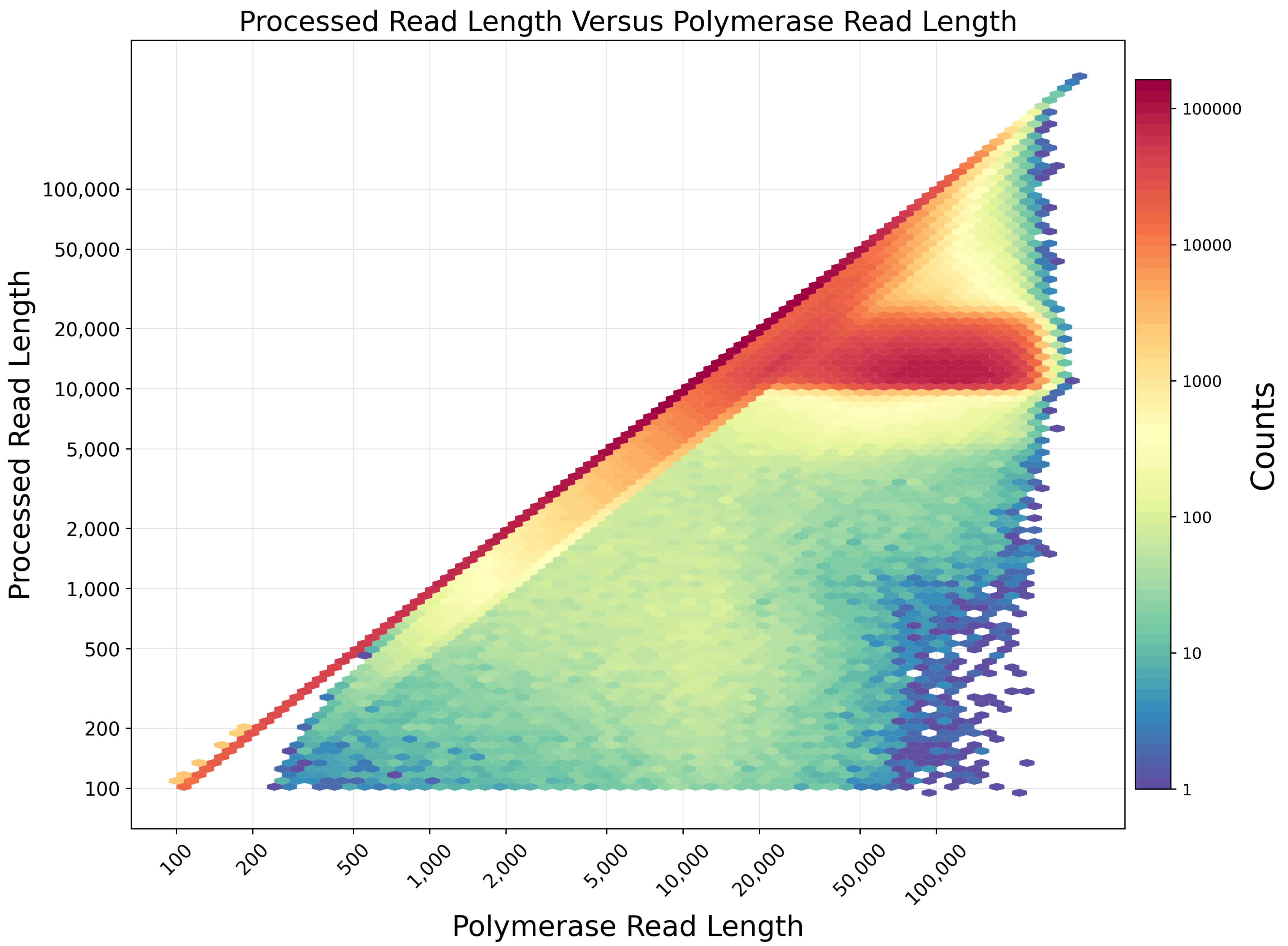

**Base Yield Density**

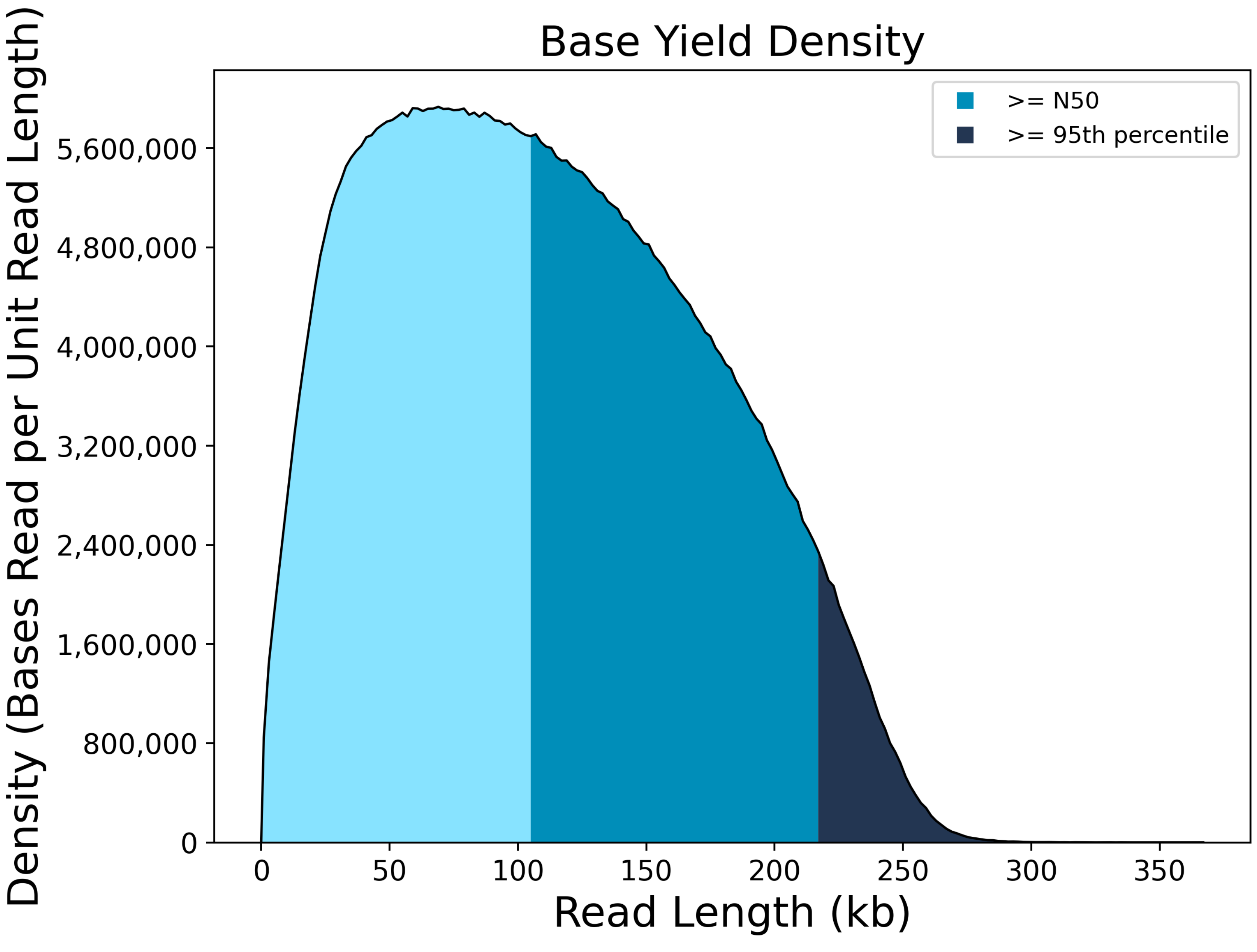

### Barcodes

#### Summary

| | |
|---|---|
| **Unique Barcodes** | 1 |
| **Barcoded HiFi Reads** | 4,995,450 |
| **Unbarcoded HiFi Reads** | 199,011 |
| **Barcoded HiFi Reads (%)** | 0.961687844032326 |
| **Barcoded HiFi Yield (bp)** | 68,367,713,586 |
| **Unbarcoded HiFi Yield (bp)** | 2,719,503,570 |
| **Barcoded HiFi Yield (%)** | 0.9617441267389595 |
| **Mean HiFi Reads per Barcode** | 4,995,450 |
| **Max. HiFi Reads per Barcode** | 4,995,450 |
| **Min. HiFi Reads per Barcode** | 4,995,450 |
| **Barcoded HiFi Read Length (mean, bp)** | 13,685 |
| **Unbarcoded HiFi Read Length (mean, bp)** | 13,665 |

#### Barcode Data

| Sample Name | Barcode | Barcode Quality | HiFi Reads | HiFi Read Length (mean, bp) | HiFi Read Quality (mean, QV) | HiFi Yield (bp) | Polymerase Read Length (mean, bp) | Polymerase Yield (bp) |
|---|---|---|---|---|---|---|---|---|
| Lep 89533 | default--default | 93.0 | 4995450 | 13685 | Q28 | 68367713586 | null | 0 |
| No Name | Not Barcoded | 0.0 | 199011 | 13665 | Q24 | 2719503570 | null | 0 |

**Number Of Reads Per Barcode**

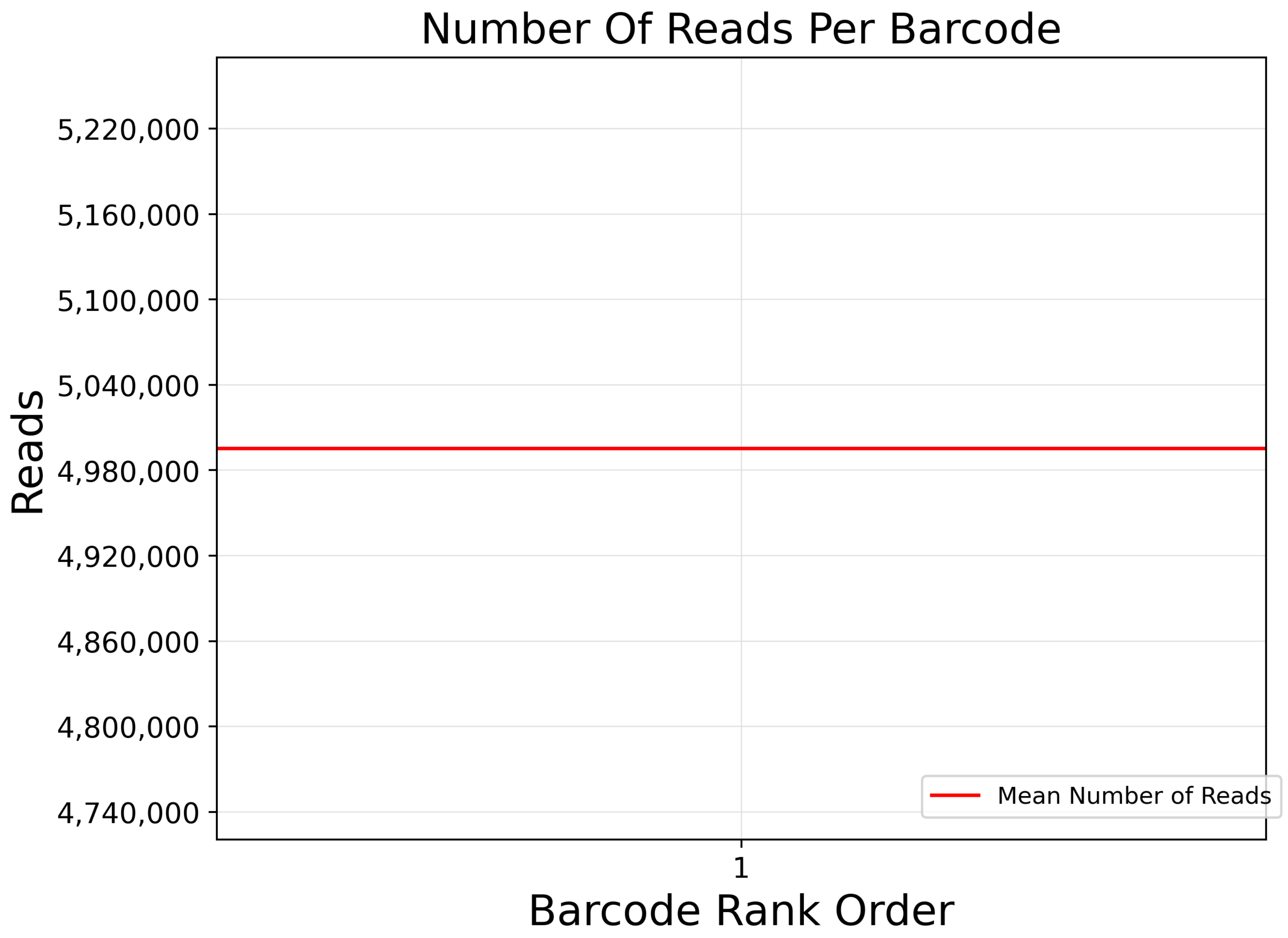

#### Barcode Frequency Distribution

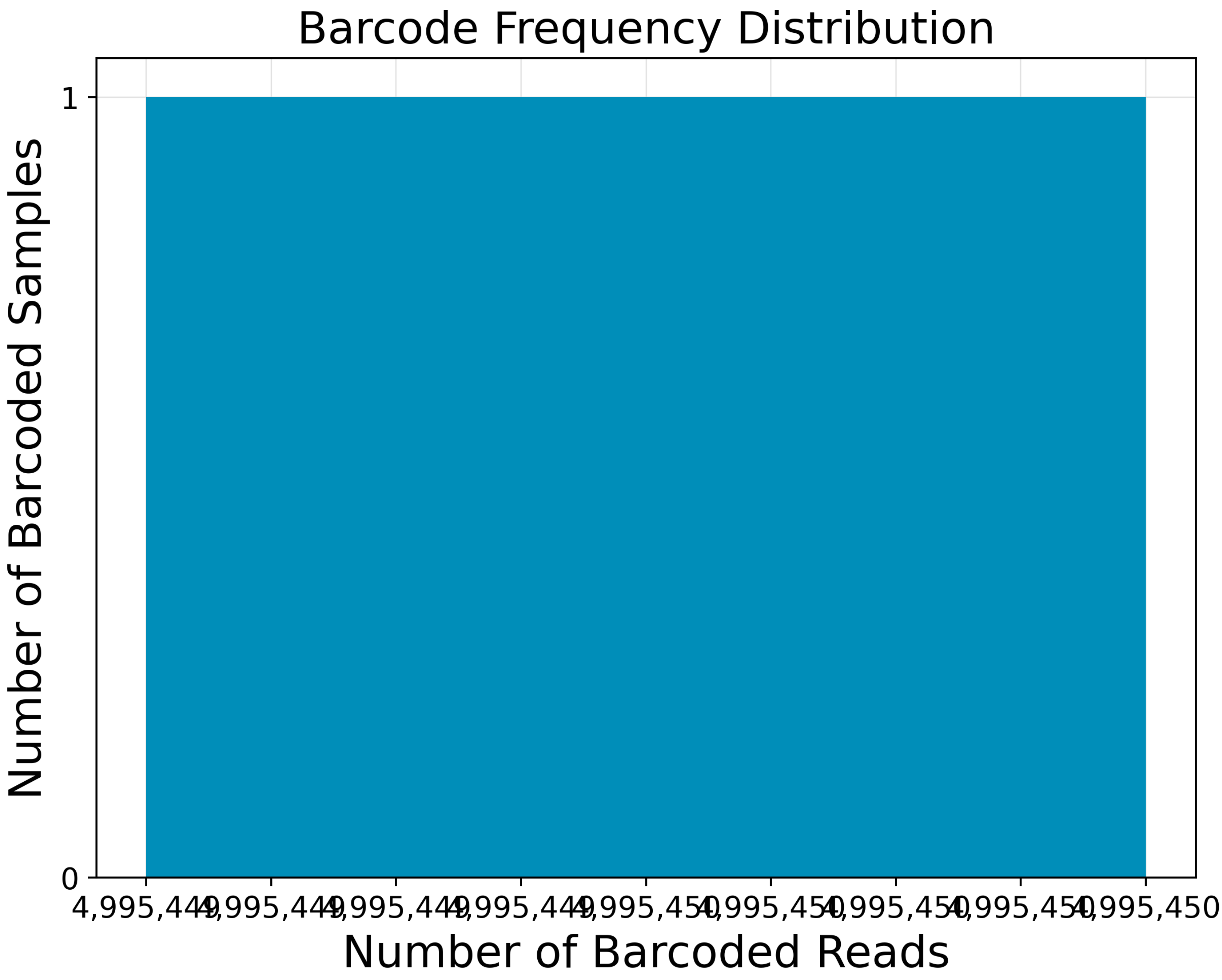
Barcode Frequency Distribution
Number of Barcoded Samples
Number of Barcoded Reads
1
0
4,995,449
4,995,449
4,995,449
4,995,449
4,995,450
4,995,450
4,995,450
4,995,450
4,995,450

#### Mean Read Length Distribution

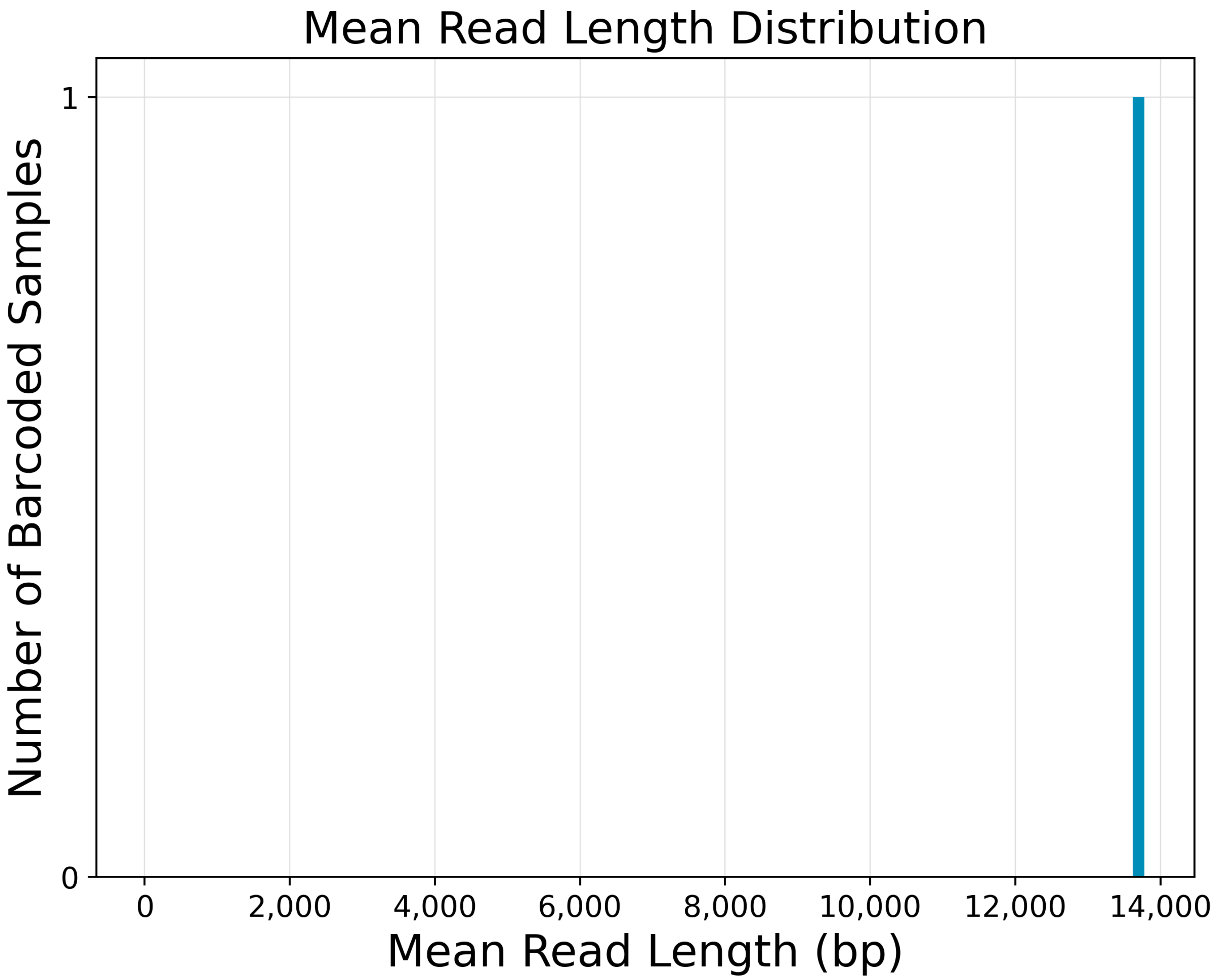

Mean Read Length Distribution
1
0
0
2,000
4,000
6,000
8,000
10,000
12,000
14,000
Mean Read Length (bp)
Number of Barcoded Samples

**Barcode Quality Score Distribution**

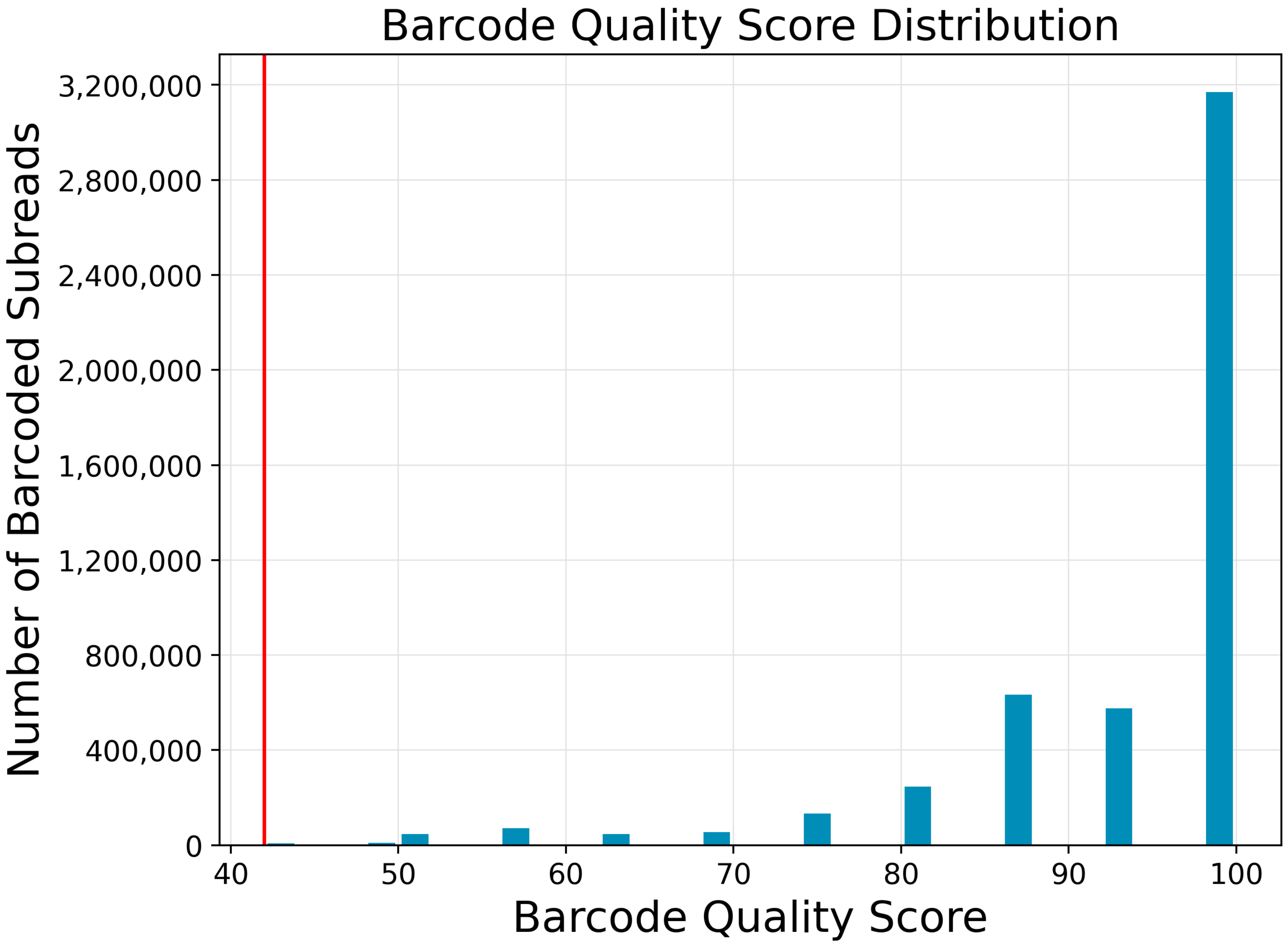

#### No Sample Setup found

### Instrument run(s)

#### Run da66caee-c666-4192-91bb-d1f902e70baf

##### Summary

| | |
|---|---|
| **Name** | Run 07.14.2023 18:18 |
| **Status** | COMPLETE |
| **Created** | 2023-07-15 00:27:39.656 |
| **Started** | 2023-07-15 00:28:50.790 |
| **Completed** | 2023-07-16 06:52:13.503 |
| **Context** | r84100_20230715_002830 |
| **Instrument Name** | 84100 |
| **Instrument Serial Number** | 84100 |
| **ICS Version** | 12.0.0.179648 |
| **Primary Analysis Version** | 12.0.0.1 |
| **Chemistry Version** | 12.0.0.172289 |

### Parent jobs (1)

### Job 5644

#### Summary

| | |
|---|---|
| **Job Type** | import-dataset |
| **Pipeline** | cromwell.workflows.sl_dataset_reports |
| **Name** | import-dataset |
| **Comments** | Description for job Import PacBio DataSet |
| **Created At** | 2023-07-16 14:30:20.353 |
| **SMRT Link Version** | 12.0.0.177059 |

#### No child jobs found
